# Ena/VASP proteins at the crossroads of actin nucleation pathways in dendritic cell migration

**DOI:** 10.1101/2022.07.29.501881

**Authors:** Sai Prasanna Visweshwaran, Hafiza Nayab, Lennart Hoffmann, Marine Gil, Fan Liu, Ronald Kühne, Tanja Maritzen

## Abstract

As sentinels of our immune system dendritic cells (DCs) rely on efficient cell migration for patrolling peripheral tissues and delivering sampled antigens to secondary lymphoid organs for the activation of T-cells. Dynamic actin polymerization is key to their macropinocytic and migratory properties. Both major actin nucleation machineries, formins and the Arp2/3 complex, are critical for different aspects of DC functionality, by driving the generation of linear respectively branched actin filaments. However, the importance of a third group of actin nucleators, the Ena/VASP family, has not been addressed yet. Here, we show that the two family members Evl and VASP are expressed in murine DCs and that their loss negatively affects DC macropinocytosis, spreading, and migration. Our interactome analysis reveals Ena/VASP proteins to be ideally positioned for orchestrating the different actin nucleation pathways by binding to the formin mDia1 as well as to the WAVE regulatory complex, a stimulator of Arp2/3. In fact, Evl/VASP deficient murine DCs are more vulnerable to inhibition of Arp2/3 demonstrating that Ena/VASP proteins contribute to the robustness and efficiency of DC migration.

## Introduction

The mammalian innate and adaptive immune defense critically relies on highly migratory leukocytes. Among those, dendritic cells (DCs) constitute the first line of defense by patrolling peripheral tissues and sampling pathogens. As professional antigen-presenting cells, immature DCs employ endocytic processes, mostly macropinocytosis, to take up antigens (1) which they process further to generate MHCII-peptide complexes for presentation on their surface. Once activated by pathogenic danger signals such as lipopolysaccharides (LPS, endotoxins that are found in the outer membrane of Gram-negative bacteria), DCs undergo dramatic alterations. This process is called maturation and enables the mature DCs to directionally migrate from the antigen acquisition site to the nearest secondary lymphoid organ to present the sampled antigen to antigen-specific naïve T-cells (2). In this manner DCs stimulate the adaptive immune response, thus acting as messengers between the innate and the adaptive immune system.

As part of the dramatic changes that are elicited by danger signals, DCs downregulate macropinocytosis, while upregulating the surface levels of MHCII and co-stimulatory molecules for interaction with T-cells. In addition, they switch to a highly directional migratory mode which includes the increased expression of the chemokine receptor CCR7 which directs them to the draining lymph node by responding to chemokines such as CCL21 and CCL19 (2). In addition, actin dynamics are markedly altered upon maturation to foster a dynamic and directed migration mode (3). It has become increasingly clear that DCs can adopt distinct migration modes triggered by the geometry of the environment (3) or internal cues (4). Especially in confined environments, DCs migrate in an amoeboid manner without requiring adhesions (5). They move by virtue of rapid cycles of actin polymerization and actomyosin-based contraction. While the importance of a protrusive actin flow for DC migration is firmly established (6), the exact function of the different actin regulatory proteins is still under debate. Earlier papers identified Rac1/2 (7), Cdc42 (8) and WASP (9), which trigger Arp2/3 mediated formation of branched actin networks, as crucial factors for DC migration. In addition, a more recent study provided evidence for a critical role of the WAVE regulatory complex (WRC), an Arp2/3 activator important for lamellipodia formation, in shaping DC migration. Deletion of a WRC subunit caused DCs to migrate with increased speed and enormous directional persistence, while they were unable to turn efficiently towards chemotactic gradients (10). Another study employing 1D confined micro-channels reported that Cdc42 and Arp2/3 slow DC motility by limiting the actin availability for a fast migration mode that is driven by RhoA and by the linear actin filament nucleator mDia1 (3) which modulate an actin pool at the cell rear.

While the importance of the actin nucleators Arp2/3 and mDia1 for DC functionality is evident, the role of a third major class of actin nucleators, the Ena/VASP protein family, has remained elusive. In fibroblasts and cancer cells the Ena/VASP protein family (comprising Mena, VASP, and Evl) is established as an important complementary orchestrator of actin filament assembly which greatly influences cell motility by binding actin as well as a variety of focal adhesion proteins and actin regulators (11,12). Also in neurons (13), T-cells (14) and leukocytes (15), Ena/VASP proteins have been shown to play crucial roles in maintaining cellular functionality.

The proteins of the Ena/VASP family are actin polymerases in their own right like formins (e.g. mDia1) and the Arp2/3 complex (16–19). However, they are also suggested to act as interaction hubs for the coordination of focal adhesion and actin regulatory proteins (20). They achieve this regulatory hub function via their actin filament binding ability in combination with their evolutionarily conserved EVH1 domain that binds to numerous interactors, including subunits of the Arp2/3 activator WRC (21). In addition, they were suggested to interact with mDia1 (22). Therefore, Ena/VASP proteins appear ideally positioned to fine-tune actin dynamics and thus migration by regulating linear actin formation via mDia1 and branched actin formation via Arp2/3. However, their relevance for DC migration has not been addressed yet.

In this study, we show that the Ena/VASP family members Evl and VASP are expressed in immature and mature DCs and that their loss affects macropinocytosis, spreading, and migration of DCs. Moreover, inhibiting Arp2/3 has more debilitating effects on the migration of Evl/VASP double knockout (DKO) DCs than on control DCs. In line with this, our interactome analysis demonstrated that also in DCs Ena/VASP proteins are connected to Arp2/3 via the WRC. In addition, we confirmed mDia1 also for this cell type as an Evl/VASP interactor. Thus, our data suggest that DCs employ three different actin nucleation machineries to shape actin dynamics for efficient migration and highlight Ena/VASP proteins as important interaction hubs at the crossroads of the different actin nucleation pathways whose loss profoundly impairs DC migration.

## Results

### Normal differentiation and maturation of Evl/VASP DKO DCs

The Ena/VASP family comprises the proteins Mena, Evl and VASP in mammals. Deletion of all three family members is embryonically lethal (13). Immune cells such as T-cells were previously shown not to express Mena (14) even though Mena is quite broadly expressed in diverse cell types ranging from neurons to cancer cells. Therefore, we decided to obtain Evl/VASP DKO mice (generously provided by Frank Gertler (13)) for our study of the role of Ena/VASP proteins in DCs. We generated bone marrow-derived dendritic cells (henceforth called DCs) from adult control and Evl/VASP DKO mice. To analyze the expression pattern of the Ena/VASP family in DCs, we probed DC lysates with specific antibodies against the three family members. We found VASP to be solidly expressed in WT DCs, and also detected Evl, albeit more faintly (Figure S1A). However, in line with expectations, we could not detect Mena with the available antibodies (Figure S1B). Neither Evl nor VASP were detected in DKO DC lysate confirming the deletion of Evl and VASP in the DKO and the specificity of the used antibodies. We also did not detect Mena in the DKO DC lysate arguing against a compensatory upregulation of this family member in DCs upon deletion of Evl and VASP (Figure S1B). This is in line with results from Evl/VASP DKO T-cells which also do not display any compensatory upregulation of Mena (14).

We ascertained the efficiency of in vitro DC differentiation by flow cytometry-based quantification of the established DC surface marker CD11c. There was no significant difference in the propensity of Evl/VASP DKO bone marrow cells to differentiate into DCs (Figure S2A,B). In addition, control and DKO immature DCs could both be efficiently converted into mature DCs (mDCs) by LPS treatment based on upregulation of the mDC surface markers CD40 and CD86 (Figure S2C-F). Thus, in vitro DC differentiation and maturation do not require Evl and VASP. This is in line with previous results from DCs that are deficient in other actin regulatory proteins such as Eps8 (23), Cdc42 (8), Hem1 (a member of the WRC) (10), WASP (9) and mDia1 (24) which all showed normal differentiation and maturation.

### Survey of focal adhesion and actin regulatory proteins in Evl/VASP DKO DCs

Since Evl and VASP constitute interaction scaffolds for focal adhesion and actin regulatory proteins, we analyzed the levels and localization of important members of these two functional groups in WT and Evl/VASP DKO iDCs and mDCs (Figures S3-S5). Overall, the levels and localization of the tested actin nucleation promoting factor WAVE2, the Arp2/3 subunit p34/ARPC2 and the formin mDia1 as well as the analyzed focal adhesion proteins vinculin and zyxin were unaltered in Evl/VASP DKO DCs (except for a very heterogenous decrease in zyxin immunofluorescence in iDCs). Thus, the loss of Evl/VASP does not impair the expression and localization of other important actin regulators.

### Decreased F-actin levels

In line with the normal expression and distribution of the major DC actin nucleators in Evl/VASP DKOs (Figures S3-S5), we did not observe any obvious differences in the total actin levels as detected by immunoblotting or in the overall localization and intensity of F-actin based on immunofluorescence stainings in DKO iDCs or mDCs (Figure 1A-C). However, quantitative analysis of total F-actin levels as measured by flow cytometry of phalloidin-labeled DCs revealed a small, but significant decrease in the amount of F-actin in DKO mDCs (Figure 1D). Thus, Evl/VASP contribute either directly by their nucleation capacity or indirectly via their impact on other actin nucleators to the generation of actin filaments in mDCs.

**Figure 1.**
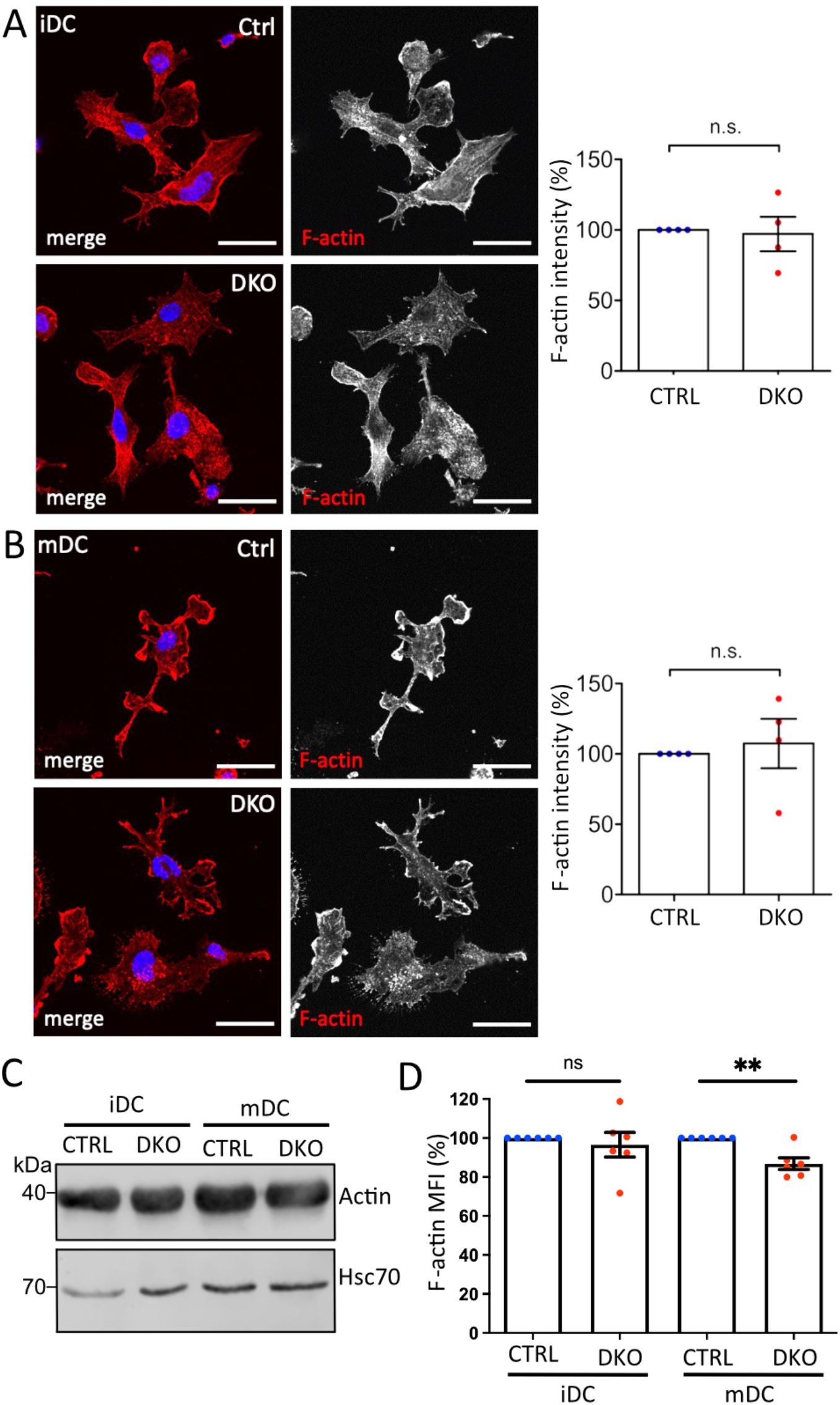
Loss of Evl and VASP results in a small reduction in the level of F-actin. (A-B) No detectable difference in F-actin localization and level by immunofluorescence. Immature (A) and mature (B) control and DKO DCs were processed for immunofluorescence and incubated with phalloidin to label F-actin. Nuclei were stained with DAPI and are depicted in blue in the merged images. Scale bar: 25 µm. Fluorescence intensities were quantified and are expressed as % of control levels (depicted as mean±SEM; statistical analysis by One-sample t-test; N=4 independent experiments; ns=non significant). (C) No difference in total actin levels. Lysates from control and DKO mDCs and iDCs were immunoblotted and probed with actin-specific antibodies to reveal total actin levels and with Hsc70-specific antibodies to control for loading. (D) Small increase in F-actin in mature DKO cells detected via flow cytometry. Immature and mature control and DKO DCs were analyzed by flow cytometry after phalloidin staining to label F-actin. Mean fluorescence intensity (MFI) is depicted as % of the respective control (depicted as mean±SEM; statistical analysis by One sample t-test; N=6 independent experiments; **p<0.01; ns=non significant).

### Impaired macropinocytosis in Evl/VASP DKO DCs

Since actin nucleation is not only involved in DC migration, but also crucial for the ability of iDCs to perform macropinocytosis, we started our functional characterization of Evl/VASP deficient DCs by analyzing their propensity to macropinocytose. To do so, we incubated control and DKO iDCs and mDCs with fluorescent FITC-dextran at 37°C to allow for macropinocytosis of the dextran. As described previously, iDCs were much more efficient at macropinocytosis than mDCs whose uptake was not much greater than that of control cells that had been incubated at 4°C in the presence of FITC-dextran to judge the contribution of background binding (Figure 2A). More importantly, the macropinocytosis of iDCs was severely reduced in absence of Evl/VASP (Figure 2A) arguing for a role of the Ena/VASP family in this actin-dependent endocytic process.

**Figure 2.**
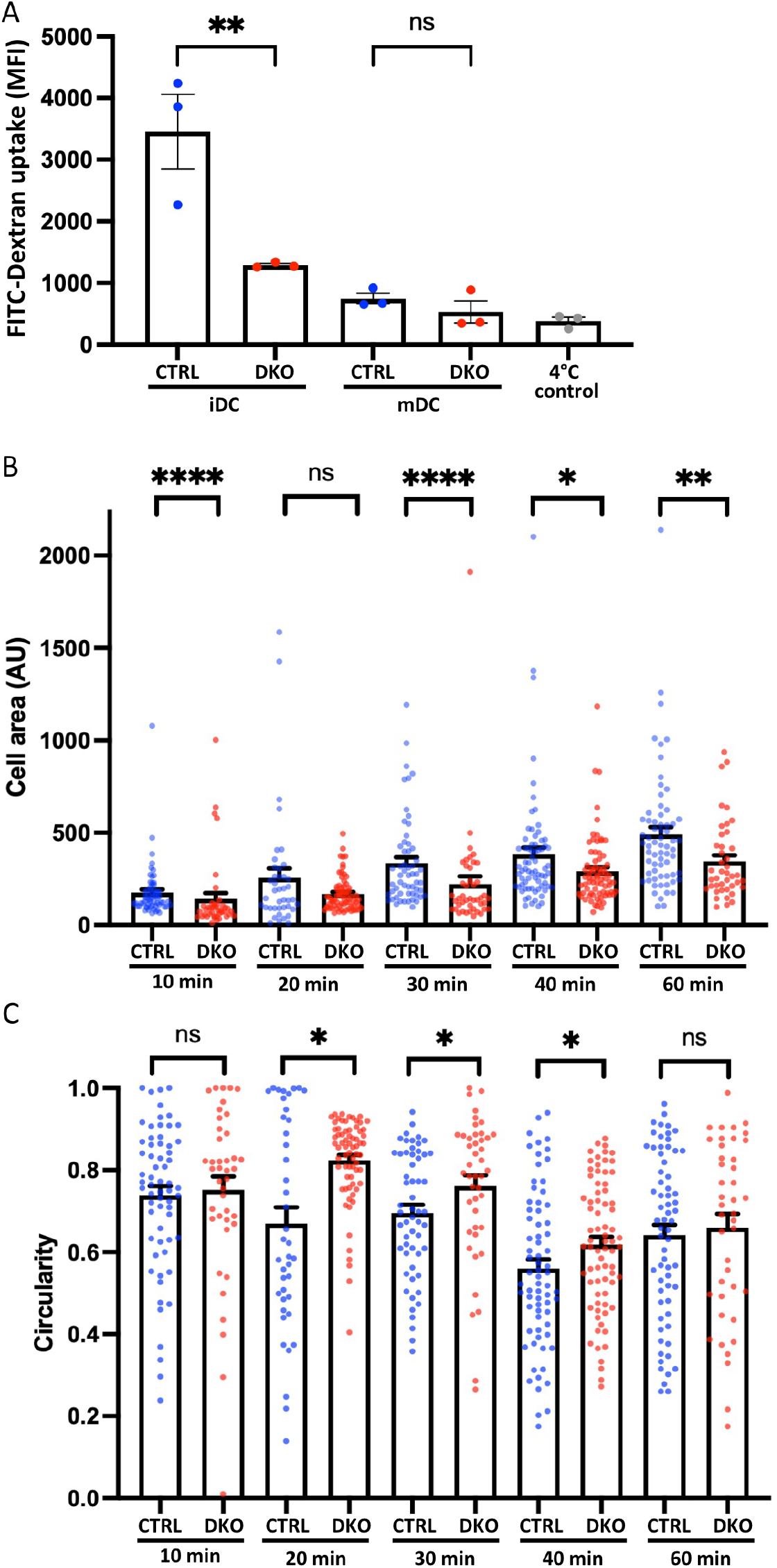
Loss of Evl and VASP impairs DC macropinocytosis and spreading. (A) Evl/VASP DKO iDCs show impaired macropinocytosis. Immature and mature control and DKO DCs were incubated with FITC-dextran for 1 h at 37°C. FITC-dextran uptake was quantified by flow cytometry. In grey: control cells left at 4°C to reveal background binding. Mean fluorescence intensity (MFI) depicted as mean±SEM (N=3 independent experiments, statistical analysis by One-Way-ANOVA followed by Tukey post-test; **p<0.01; ns=non significant). (B,C) Mature control and DKO DCs were allowed to spread on fibronectin-coated cover slips for the indicated time points. Cells were processed for immunofluorescence and incubated with phalloidin to stain F-actin and thus visualize cell shape. Cell area (B; in arbitrary units, AU) and circularity (C) were quantified based on Image J (depicted as mean±SEM; statistical analysis by Mann-Whitney test; N=41-74 cells from 3 independent cultures processed in 2 independent experiments; *p<0.05; **p<0.01; ***p<0.001; ****p<0.0001; ns=non significant).

### Impaired spreading of Evl/VASP DKO DCs

Due to the known links of Evl/VASP proteins to focal adhesions, we decided to evaluate the impact of their loss on DC spreading. For that, we seeded mDCs on fibronectin-coated glass coverslips and allowed them to spread for different amounts of time before rapid fixation. Fixed cells were stained with phalloidin to easily recognize cell boundaries for the quantification of cell area and cell shape. Using the increase in cell area as a readout for cell spreading, we detected already 10 min after cell seeding a significant decrease in the spreading of DKO DCs, which grew more pronounced at 30 to 60 min post-seeding (Figure 2B). In parallel, we analyzed the circularity of cells to track the symmetry breaking which usually accompanies cell spreading. While control and DKO DCs both started out with a high circularity index, the circularity of the control cells started soon to decrease. However, the DKO DC population continued to display a high circularity index for a long time, which only clearly decreased after 40 min suggesting a delay in the symmetry breaking (Figure 2C).

### Altered migration in 2D

The impact of Ena/VASP family proteins on cell migration has so far most extensively been analyzed in the context of mesenchymal migration on 2D surfaces. Therefore, we started our analysis of Evl/VASP DKO DC motility by seeding mDCs on fibronectin-coated cell culture dishes and tracking their movement in the presence of the chemokine CCL19 over several hours (Figure 3A). In line with reports for mesenchymal cells such as fibroblasts (12), the loss of Evl and VASP significantly impaired the Mean Squared Displacement (MSD) of the DKO DCs (Figure 3B), which is a measure for the area cells explore over a given time period (25,26). This reduction in MSD is a direct consequence of a significant decrease in DKO speed (Figure 3C) and directionality (Figure 3D).

**Figure 3.**
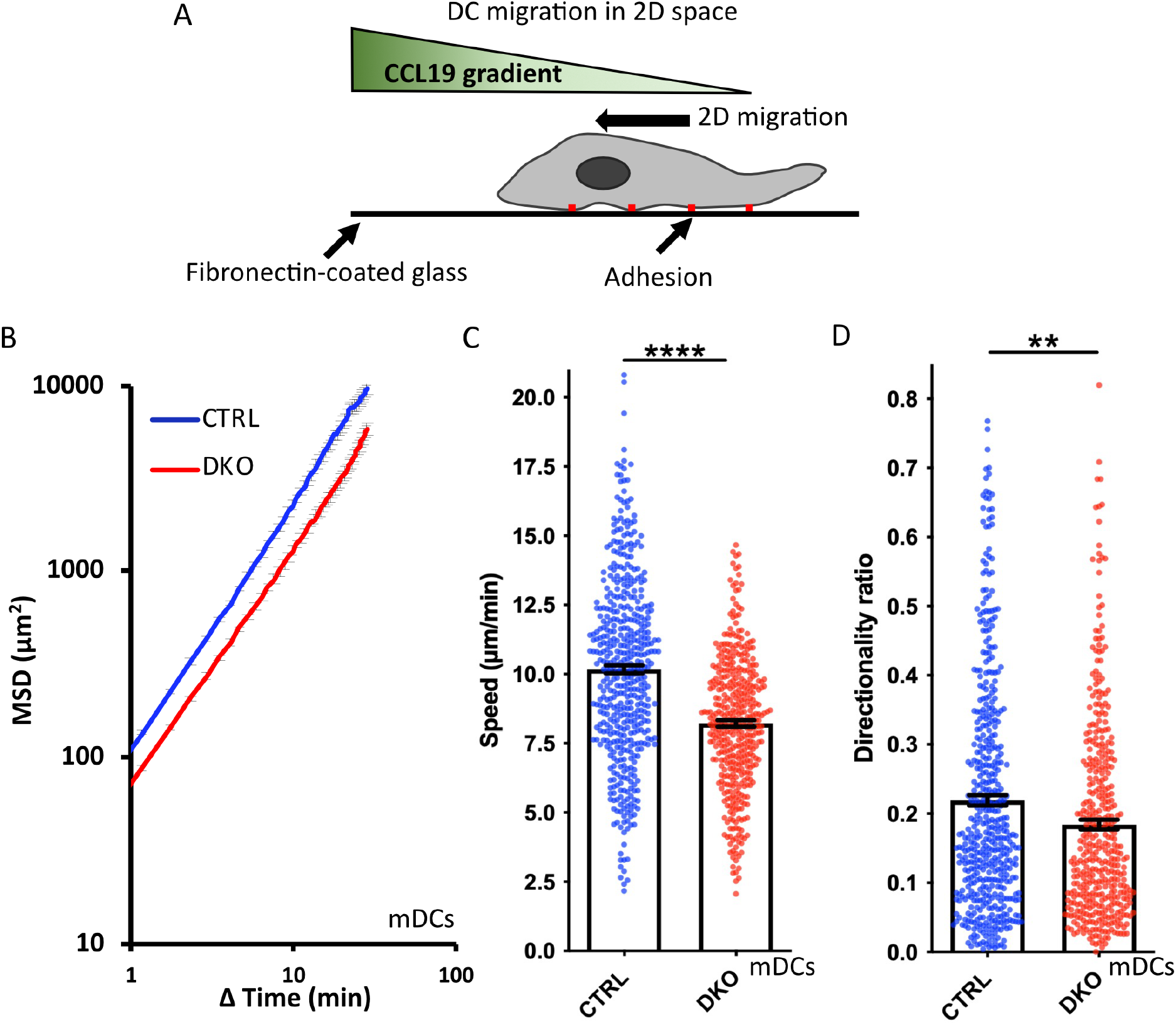
Loss of Evl and VASP impairs DC chemotaxis in 2D. (A) Illustration of experimental setup. (B-D) Mature control and DKO DCs were seeded into a migration chamber. DCs were allowed to migrate for 6 h towards the chemokine CCL19. Their migratory behavior was evaluated based on bright-field images obtained at 35 s intervals. (B) Quantification of Mean Squared Displacement (MSD). Speed (C) and directionality (D) are depicted as mean±SEM (N(control)=493 and N(DKO)=420 cells from 3 independent experiments; statistical analysis by Mann-Whitney test; **p<0.01; ****p<0.0001; ns=non significant).

### Impaired migration in confined environments

As discussed in the introduction, DC migration in confined environments differs substantially from their migration on 2D surfaces. The migration mode becomes independent of integrin-based adhesions and largely relies on actin polymerization cycles. In line with this, several actin regulatory proteins were reported to have much stronger effects on DC migration in confined environments than in 2D (8,23). Therefore, we went on to assess the migration of Evl/VASP DKO DCs in the under-agarose assay in presence of CCL19. In this assay, cells are partially confined by being squeezed between a layer of agarose and a coverslip (Figure 4A). Since iDCs and mDCs are reported to rely to varying extents on the different actin nucleation machineries (3), we tested both types of DCs in this assay. For both kinds of DCs, the speed of migration was decreased in absence of Evl and VASP like in the 2D migration assay, however, more strongly for mDCs (Figure 4C,F). This was also clearly reflected in the much more pronounced reduction in MSD for the mDCs (Figure 4B,E). Interestingly, in contrast to the results from the 2D chemotaxis assay, directionality was not decreased, but unaffected in the case of the DKO iDCs and even slightly increased for the DKO mDCs (Figure 4D,G).

**Figure 4.**
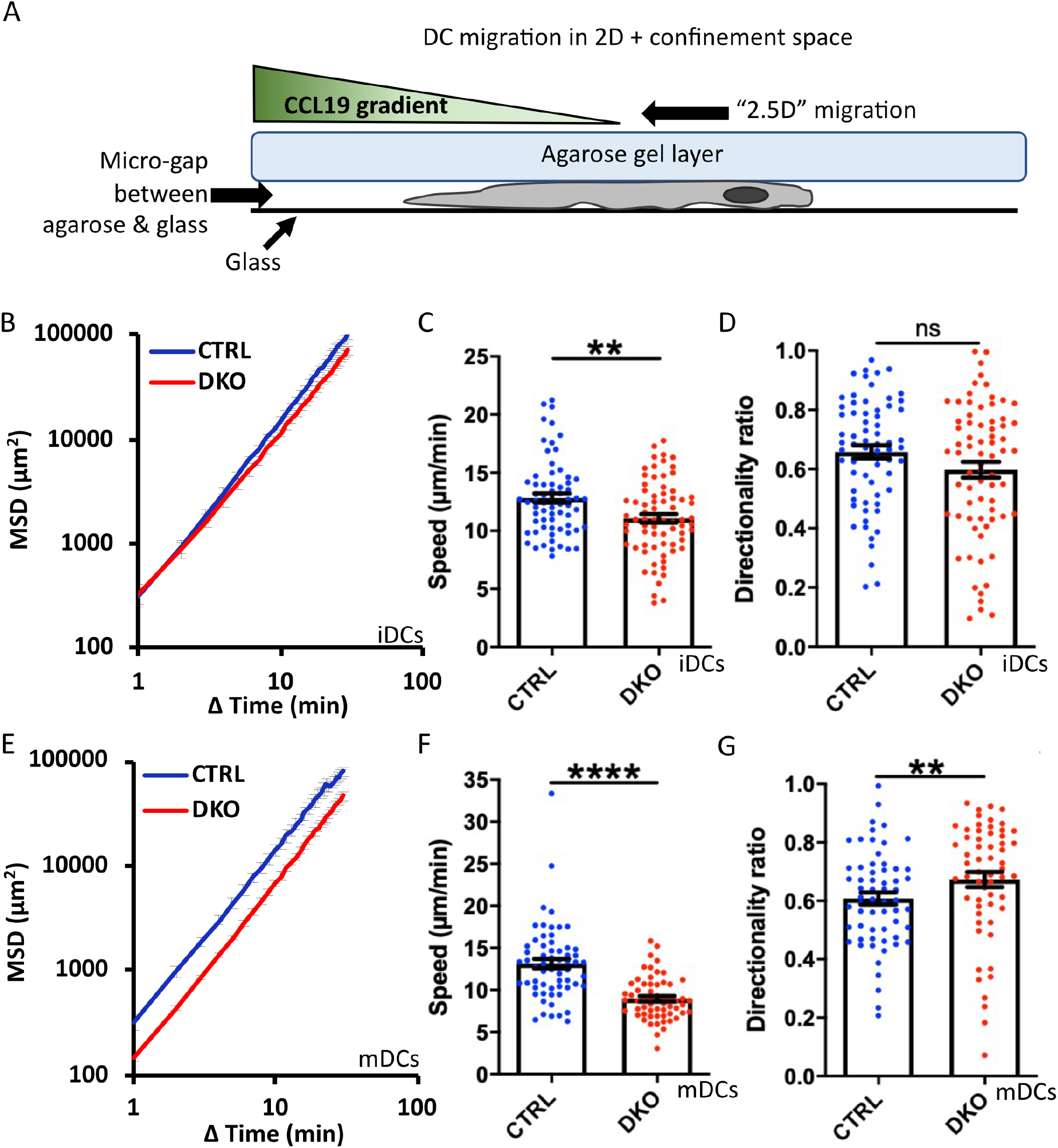
Loss of Evl and VASP impairs DC migration in confined environments. (A) Illustration of experimental setup. Immature (B-D) or mature (E-G) control and DKO DCs were allowed to migrate under agarose for 6 h attracted by CCL19. Their migratory behavior was evaluated based on bright-field images obtained at 1 min intervals. (B,E) Quantification of Mean Squared Displacement (MSD). Speed (C,F) and directionality (D,G) are depicted as mean±SEM (N(iDC control)=67, N(iDC DKO)=72, N(mDC control)=61, N(mDC DKO)=59 cells from 3 independent experiments; statistical analysis by Mann-Whitney test; **p<0.01; ****p<0.0001; ns=non significant).

To finally study the migration of Evl/VASP DKO DCs in a more complex 3D environment, we turned to collagen gels which are widely used to mimic the collagen-rich tissue DCs encounter *in vivo*. Here, differences between DKO iDCs and mDCs were most pronounced as DKO mDCs displayed a substantially reduced MSD whereas DKO iDCs rather gained MSD compared to control cells (Figure 5A,B,E).The loss of Evl/VASP did not affect the speed of iDCs within the collagen gel at all, while the speed of mDCs was significantly decreased (Figure 5C,F). Directionality was affected for both types of DCs upon loss of Evl/VASP, however, in opposite directions. iDCs substantially gained in directionality in absence of Evl/VASP (Figure 5D) while mDCs suffered from a loss of directionality (Figure 5G).

**Figure 5.**
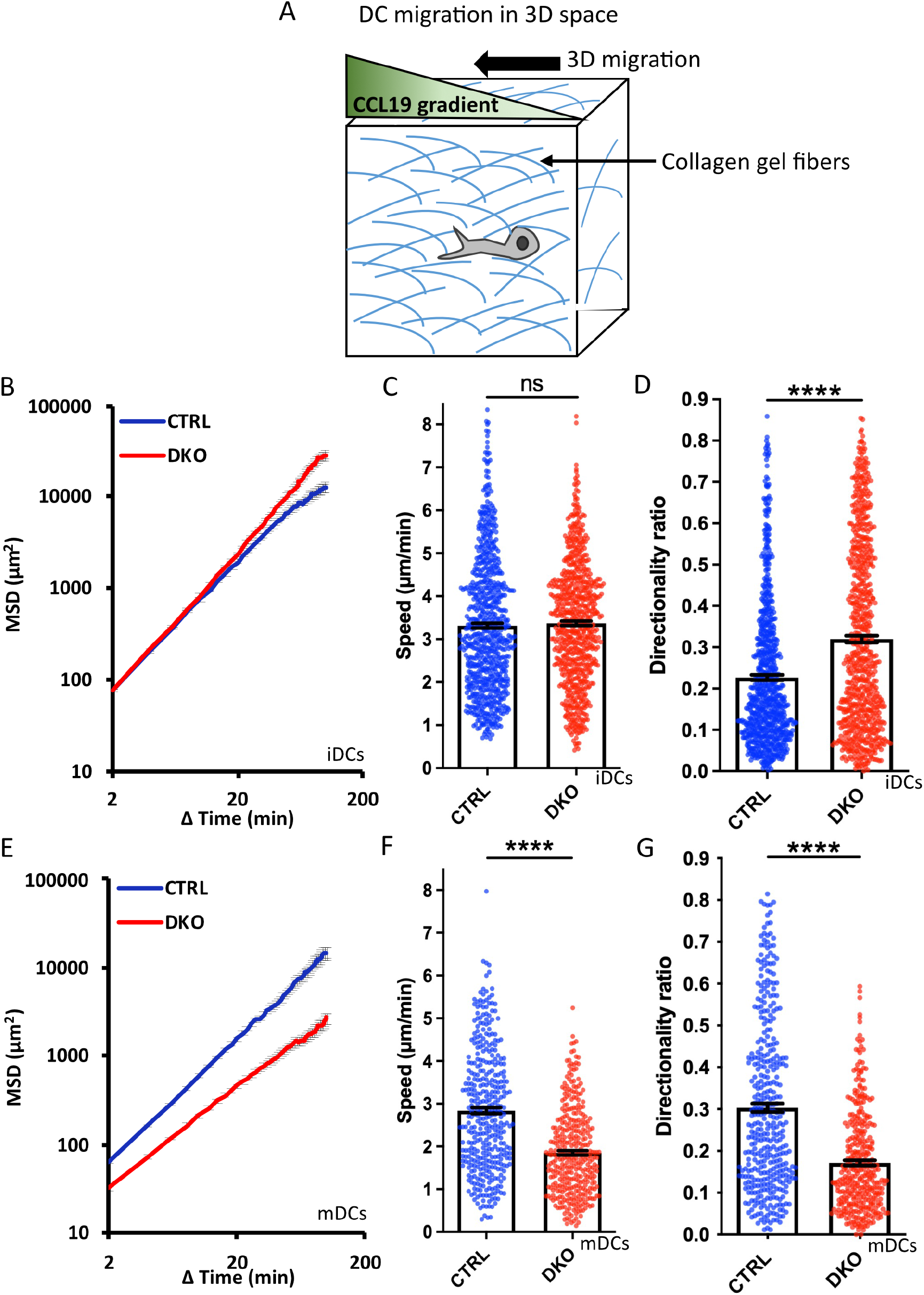
Loss of Evl and VASP alters DC migration towards CCL19 gradients in 3D collagen gels. (A) Illustration of experimental setup. Immature (B-D) or mature (E-G) control and DKO DCs were embedded into 1.9 mg/ml collagen gels. The upper gel surface was covered with medium containing 650 ng/ml CCL19. DC migration was monitored by bright-field real-time microscopy for 6 h. Images were obtained at 2 min intervals and analyzed in automated manner. (B,E) Quantification of Mean Squared Displacement (MSD). Speed (C,F) and directionality (D,G) are depicted as mean±SEM (N(iDC control)=785, N(iDC DKO)=750, N(mDC control)=389, N(mDC DKO)=372 cells from 6 (for iDCs) respectively 3 (for mDCs) independent experiments; statistical analysis by Mann-Whitney test; ****p<0.0001; ns=non significant).

We used the marked impairments of DKO mDCs in collagen gels to also address the question of the respective contributions of Evl and VASP to the observed phenotypes. For this, we tested all possible genotype combinations (Evl^wt^/VASP^wt^; Evl^ko^/VASP^wt^; Evl^wt^/VASP^ko^; Evl^ko^/VASP^ko^) separately for their 3D migration abilities. Speed and directionality only showed substantial impairments in the DKO mDCs revealing that Evl and VASP both play a role in DC migration and likely compensate for the loss of each other (Figure S6). This is reminiscent of recent data from fibroblasts where Evl also appears to be expressed at low levels, but its additional elimination still had a clear impact on migration (12).

### Evl/VASP as connecting scaffold between actin nucleation pathways

The use of different actin polymerization machineries during DC migration raises the question of their inter-dependence and spatio-temporal coordination. With their known large interactome, which includes many actin regulatory molecules, Ena/VASP proteins are ideally suited to act in the fine-tuning of actin dynamics. Since it has not been addressed in how far the established interactome of Ena/VASP proteins is recapitulated in DCs and whether DC maturation leads to differences in their interaction profile, we decided to investigate Ena/VASP interactions in DCs by an unbiased approach.

Ena/VASP proteins have a tripartite structure. The C-terminal EVH2 domain contains G- and F-actin binding sites and enables tetramerization. The central Pro-rich sequence is known to bind to several SH3- and WW-domain containing proteins. However, by far the most interactions target the N-terminal EVH1 domain which is largely similar between Evl and VASP. Therefore, we used the GST-tagged EVH1 domain of VASP, which can be easily recombinantly expressed and purified, as bait to pull down interactors from lysates of iDCs and mDCs. Canonical interactors of the EVH1 domain are known to bind via a Pro-rich so-called FPPPP motif ([D/E]FPPPxDE). To verify the specificity of the precipitated interactors, we performed control pulldowns with the GST-VASP-EVH1 domain in presence of a specific inhibitor of the FPPPP interaction site (27) reasoning that this should outcompete any interactors that employ the canonical binding mode.

Eluates from two independent experiments were analyzed by mass spectrometry-based on iDC or mDC lysates (Figure 6A,B). Of the obtained proteins we included only those as hits in Table 1 that were either significantly enriched (i.e., log2 ratio of intensities > 2.5 in both experiments) in the no-inhibitor condition or where the potential interactor was not detectable at all upon inhibitor treatment (for detailed MS data please see the supplementary excel file 1). In line with earlier reports, we detected the known interactors RIAM, lamellipodin, and zyxin, which link Ena/VASP proteins to focal adhesions and lamellipodia. In addition, we prominently identified components of the WRC, one of the nucleation promotion factors (NPFs) that stimulates the actin nucleator Arp2/3. Among the retrieved WRC components were Hem 1, Hem 2, Cyfip1, Cyfip2, Abi1, Abi3 and Wave2 indicating that DCs express multiple isoforms of the different WRC subunits. Wave 1 and Wave 2 have been shown to directly interact with Ena/VASP proteins before, however, the binding to Abi is believed to be the more important link between Ena/VASP and the WRC (21,28). We also found a subunit of the Arp2/3 complex itself. However, the Arp2/3 complex has not previously been shown to interact directly with VASP, but might be precipitated indirectly via the WRC. Finally, we confirmed the interaction between VASP and the formin mDia1 that had been suggested as an interactor in one earlier study (22). In fact, mDia1 showed the highest enrichment over the inhibitor-treated sample in our DC analysis. Using a pulldown approach as outlined above, but followed by immunoblotting, we successfully verified the interactions of the VASP-EVH1 domain with the WRC and mDia1. Thus, our data support the notion that Ena/VASP proteins are not only part of the cellular actin nucleation machinery, but also linkers between different actin nucleation pathways.

**Table 1:**
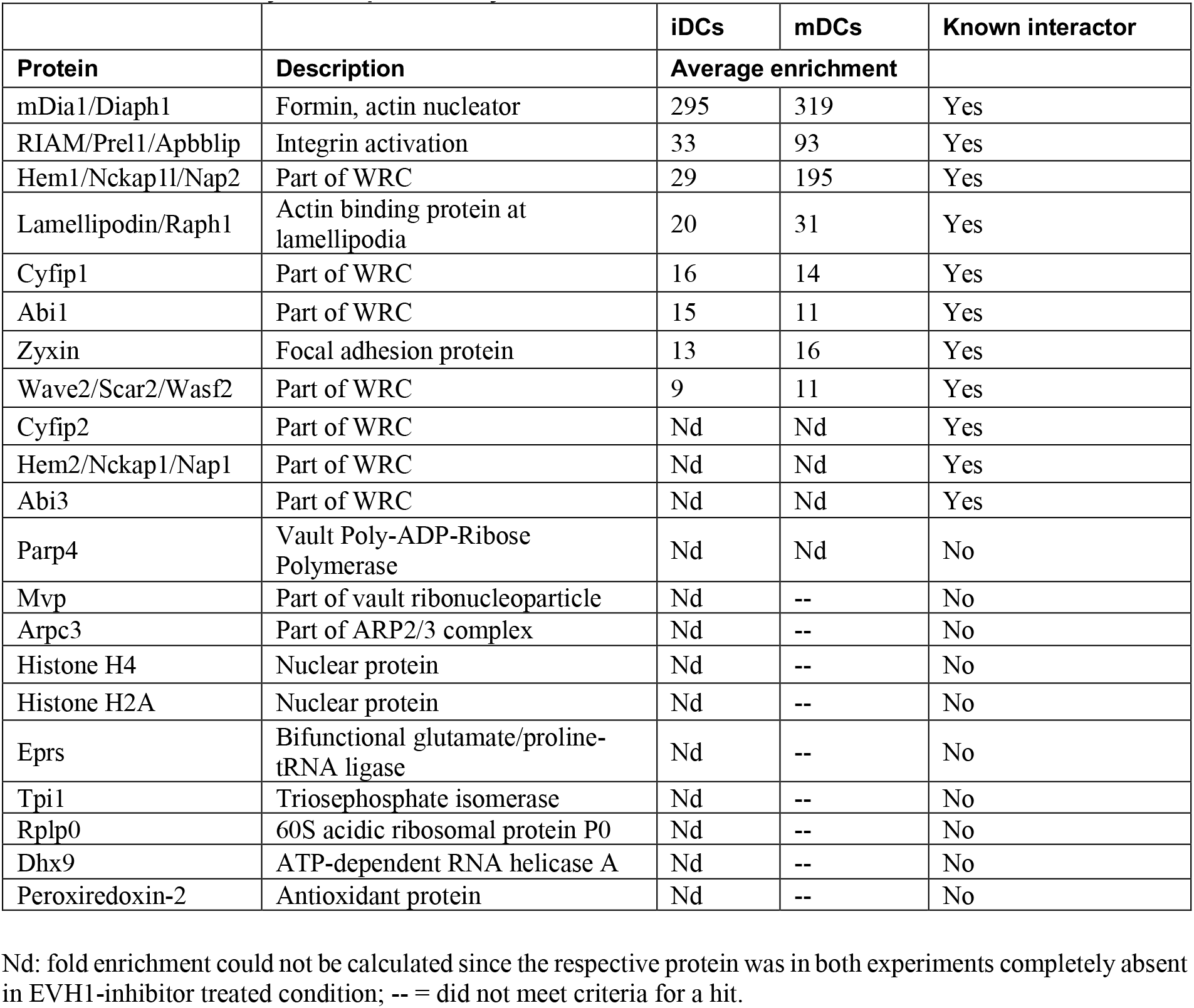
Hits obtained by mass spectrometry.

**Figure 6.**
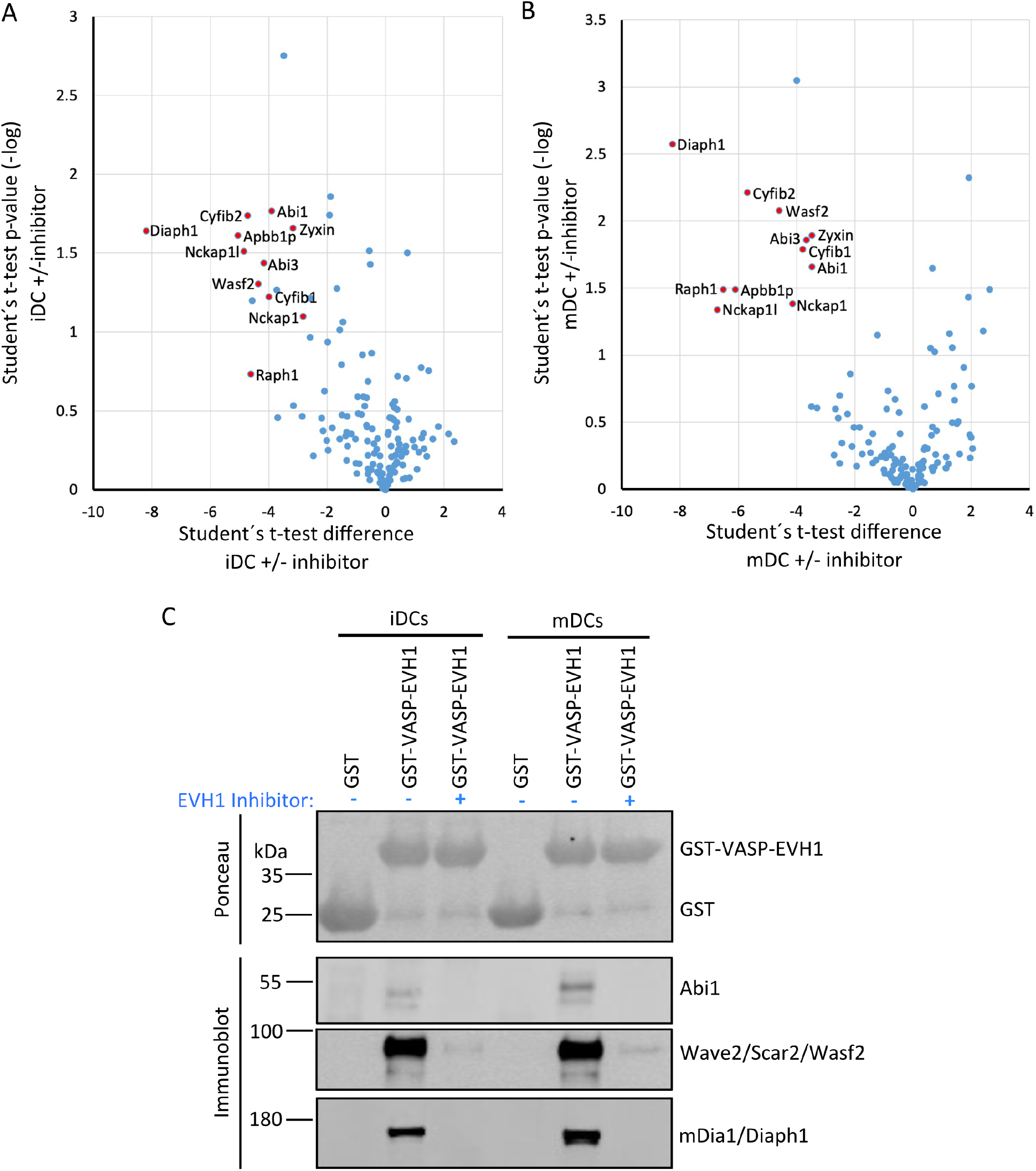
The VASP-EVH1 domain interacts with Wave2 and mDia1 in DCs. Recombinant GST-VASP-EVH1 and GST as control were purified from bacteria with GST-beads and incubated with detergent-extracted lysate of immature or mature DCs in the presence or absence of an EVH1-specific inhibitor. After thorough washing, proteins were eluted and analyzed by mass spectrometry (A,B) or by immunoblotting with the indicated antibodies (C). (A,B) Volcano plots illustrate the inhibitor-sensitive hits obtained by mass spectrometry for iDCs (A) and mDCs (B). (C) Immunoblotting confirms that Abi1 and Wave2 (as representatives of the WRC) and mDia1 from iDCs and mDCs show specific binding to the EVH1 domain of VASP which is inhibited by the EVH1 domain-specific inhibitor.

In addition to the actin dynamics related interactors, the mass spectrometry analysis also revealed several unrelated proteins that have no obvious links to the known functions of Ena/VASP family members and will need to be verified in the future.

In summary, our results show that the core interactome of the VASP-EVH1 domain is highly conserved in DCs and also does not differ between iDCs and mDCs arguing for a highly conserved function of Evl/VASP proteins in the regulation of actin dynamics across cell types, especially in the orchestration of actin nucleation pathways.

### Loss of Evl/VASP increases the vulnerability of DCs to impairments of other actin nucleation pathways

Ena/VASP proteins are actin nucleators and thus might act redundantly with other nucleation machineries. Alternatively, they might function as interaction hubs for the coordination of the different actin nucleation pathways. Therefore, we wondered whether loss of Evl and VASP renders DCs more vulnerable to the inhibition of any of the other actin nucleators. To address this question, we used the 3D migration assay and first treated control and DKO iDCs and mDCs with the established formin inhibitor SMIFH2, using a rather low concentration of 5 µM and a higher concentration of 10 µM to be in the dynamic range. Both concentrations are well below the 25 µM to 50 µM range that has been reported to lead to immediate DC rounding and partial cell death (4,10). In this context it has additionally to be taken into consideration that SMIFH2 was recently reported to have side effects on myosins. In fact, SMIFH2 was shown to inhibit human non-muscle myosin IIA activity with an IC_50_ of ~50 µM (29). Even though we used with 5 to 10 µM very low SMIFH2 concentrations, we cannot exclude that myosin IIA which plays an important role in DC migration (30) is also partially affected by the treatment.

At 5 µM SMIFH2 had already significant, but rather mild effects on control iDCs and mDCs in regards to speed and directionality. To facilitate comparisons across all samples, we normalized our results to the untreated condition of the respective genotype (Figure 7A,B,D,E,G,H,J,K). For both DC types, speed and directionality of 5 µM SMIFH2-treated control cells decreased in the range of 22-36% compared to untreated controls. In contrast to that, the inhibitor effect on Evl/VASP DKO DCs was much more pronounced. Here, the treatment led to reductions ranging from 47-74%. To more easily compare the extent of inhibitor effects on DKO DCs vs controls, we calculated the ratio of the respective reductions in speed or directionality (% reduction DKO divided by % reduction control; Figure 7C,F,I,L). This illustrates that the effect of the inhibitor on speed and directionality at 5 µM was about 1.7-to 3-fold higher in the case of DKO iDCs and mDCs than in controls.

**Figure 7.**
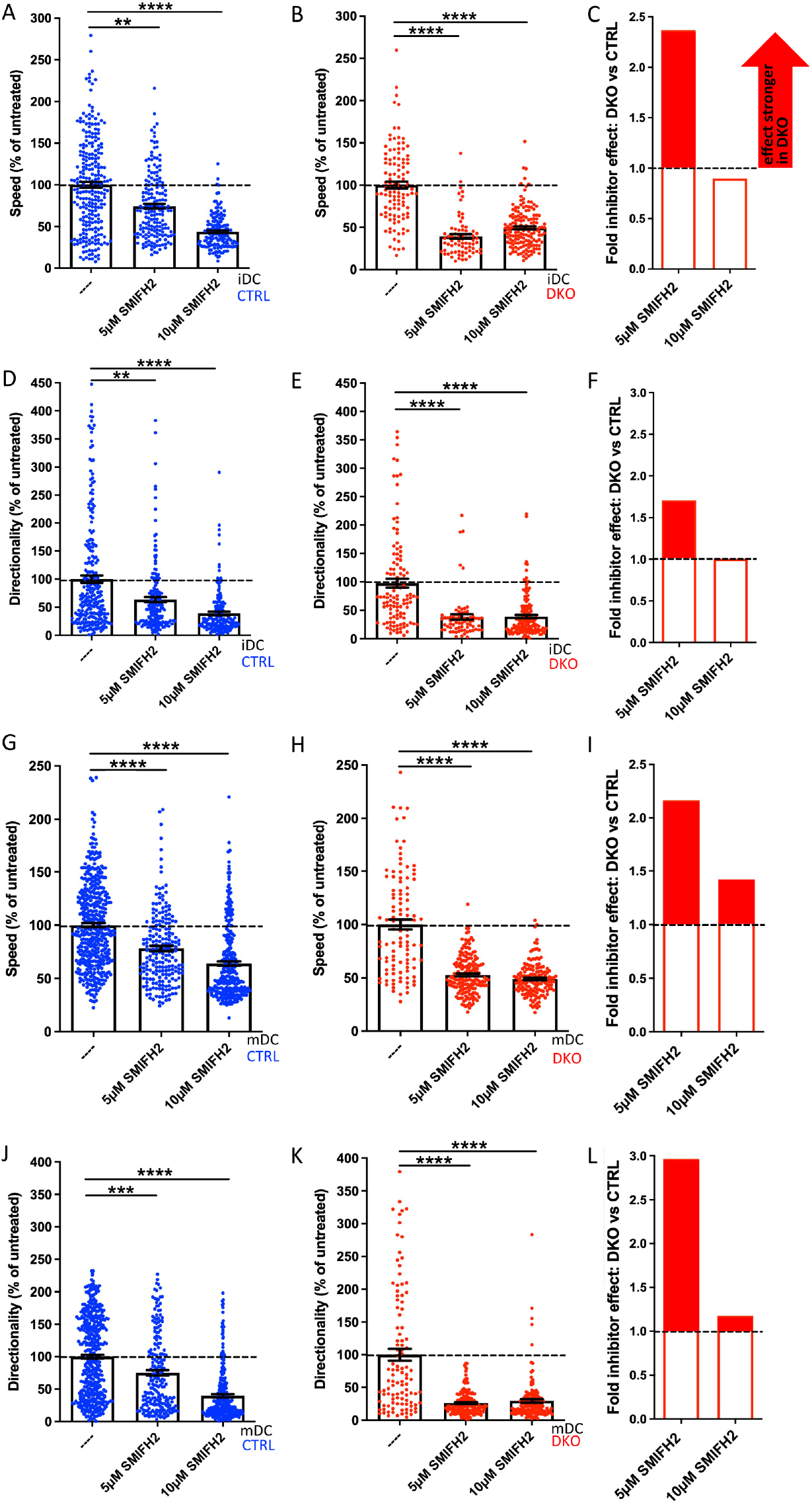
EVL/VASP DKO DCs show increased vulnerability to treatment with the formin inhibitor SMIFH2. Immature (A-F) or mature (G-L) control and DKO DCs were embedded into 1.9 mg/ml collagen gel. The upper gel surface was covered with medium containing 650 ng/ml CCL19. DCs were either left untreated, or incubated with 5 µM or 10 µM of the formin inhibitor SMIFH2 (present in gel and medium). DC migration was monitored by bright-field real-time microscopy for 6 h. Images were obtained at 2 min intervals and analyzed in automated manner. Speed (A,B,G,H) and directionality (D,E,J,K) are depicted as mean±SEM after normalization to the untreated condition of the respective genotype (N(iDCs)=72-253 cells from 2-3 independent experiments; N(mDCs)=106-438 cells from 2-4 independent experiments; statistical analysis by Kruskal-Wallis test with Dunn’s multiple comparison test on non normalized data; **p<0.01; ***p<0.001; ****p<0.0001). (C,F,I,L) Quantification of the extent of the inhibitor effect on DKO vs control DCs (i.e. the extent of the reduction in either speed (C,I) or directionality (F,L) upon inhibitor application calculated as % reduction DKO divided by % reduction control) at 5 µM or 10 µM visualizes the greater SMIFH2 effect on DKO migratory parameters, especially at 5 μM.

In case of the already strongly affected DKO iDCs and mDCs the doubling of the inhibitor concentration had no extra effect, while speed and directionality of control cells decreased further at 10 µM approximating the level that the DKO cells had reached already upon treatment with 5 µM inhibitor. Therefore, the previous difference between the genotypes largely disappeared upon the higher concentration suggesting that the inhibitor reaches its maximum potency at 10 µM and that full formin and/or myosin inhibition is incompatible with efficient cell speed and directionality. In summary, this experiment underlines that loss of Evl and VASP renders DCs highly vulnerable to even a partial inhibition of formins and potentially myosins that can still be largely tolerated by control cells.

In the second round of experiments, we tested whether the same holds true for an inhibition of the Arp2/3 complex (Figure 8). For this, we performed the 3D migration assays in the presence of the Arp2/3 inhibitor CK666 choosing 25 µM as low and 50 µM as high concentration. These concentrations are well below the 100 µM that have been used in previous studies without negative impact on DC viability (4).

**Figure 8.**
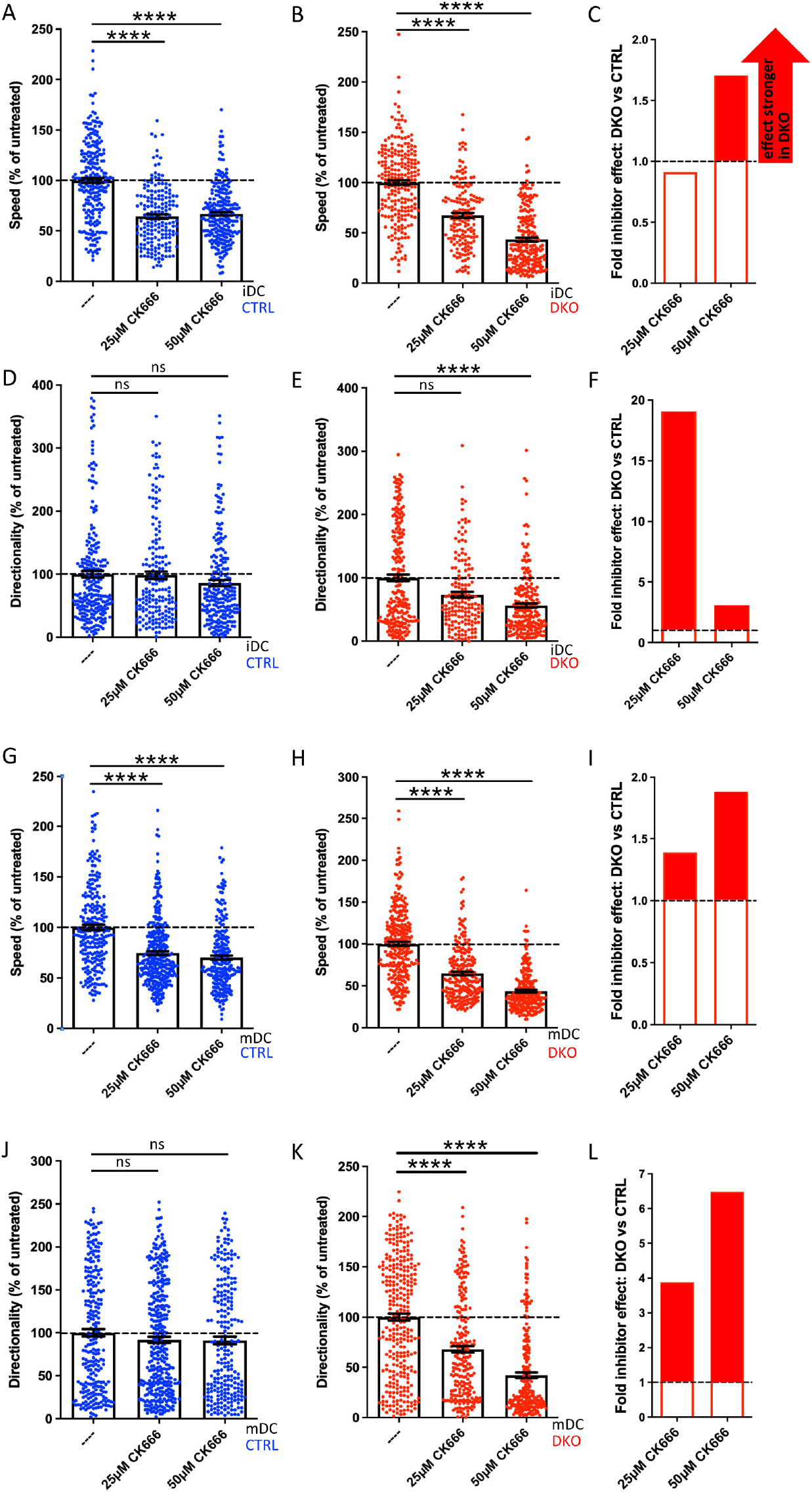
Evl/VASP DKO DCs show increased vulnerability to Arp2/3 inhibition. Immature (A-F) or mature (G-L) control and DKO DCs were embedded into 1.9 mg/ml collagen gel. The upper gel surface was covered with medium containing 650 ng/ml CCL19. DCs were either left untreated, or incubated with 25 µM or 50 µM of the Arp2/3 inhibitor CK666 (present in gel and medium). DC migration was monitored by bright-field real-time microscopy for 6 h. Images were obtained at 2 min intervals and analyzed in automated manner. Speed (A,B,G,H) and directionality (D,E,J,K) are depicted as mean±SEM after normalization to the untreated condition of the respective genotype (N(iDCs)=161-241 cells from 2 independent experiments; N(mDCs)=233-337 cells from 2-3 independent experiments; statistical analysis by Kruskal-Wallis test with Dunn’s multiple comparison test on non normalized data; ***p<0.001; ns, non significant). (C,F,I,L) Quantification of the extent of the inhibitor effect on DKO vs control DCs (i.e. the extent of the reduction in either speed (C,I) or directionality (F,L) upon inhibitor application calculated as % reduction in DKO divided by % reduction in control) at 5 µM or 10 µM visualizes the greater SMIFH2 effect on DKO migratory parameters, especially at 5 μM.

In contrast to SMIFH2 treatment, Arp2/3 inhibition of control iDCs and mDCs at both concentrations only decreased their speed by 25-36% as the normalized data reveal (Figure 8A,G), but did not significantly affect their directionality (Figure 8D,J). Again, Evl/VASP DKO iDCs and mDCs were more vulnerable to inhibitor treatment. Especially at the higher CK666 concentration, their speed was more strongly diminished than that of the control cells (Figure 8B,C,H,I) with the reductions reaching ~56% which corresponds to a 1.7-to 1.9-fold higher inhibitor effect. Even more strikingly, in contrast to the control cells, the treated DKO cells showed a significant reduction in directionality that amounted to 48-58% at the higher inhibitor concentration which corresponds to a 3-to 7-fold higher efficiency of CK666 to inhibit directionality of DKO cells as compared to its effect on control cells (Figure 8E,F,K,L).

In contrast to our experiments with SMIFH2, with the chosen concentrations of CK666 we did not reach the point where control and DKO cells showed the same extent of speed and directionality reduction. This could either be due to the possibility that Arp2/3 inhibition was not yet complete at 50 µM or it might suggest that inhibition of Arp2/3 on its own is not sufficient to further diminish speed and directionality. In summary, DCs that have lost Evl and VASP functionality are less able to tolerate inhibition of additional actin regulators.

## Discussion

Research over the past years has outlined a division of labor between the two major actin nucleation machineries in DCs. Arp2/3-dependent branched actin formation is on the one hand essential for macropinocytosis (3,8). On the other hand, like in mesenchymal cells, Arp2/3-mediated branched actin networks underlie lamellipodial protrusion (8,10). However, in confined environments, these lamellipodia do not promote adhesive migration, but rather partake in making directional choices (10). Actual translocation of the cell body appears instead to be driven mainly by formins such as mDia1 which do not contribute to macropinocytosis (3), but shape an actin pool at the cell rear for fast migration. How do Ena/VASP proteins fit into this picture?

In regards to macropinocytosis loss of Ena/VASP with its strong impairment of FITC-dextran uptake clearly resembles the loss of Arp2/3 (3) or its regulator Cdc42 (3,8). Thus, Ena/VASP proteins and the Arp2/3 complex might act within the same pathway or fulfill overlapping functions in macropinocytosis.

In regards to cell spreading, loss of Evl/VASP delays the increase in the adherent cell area of DCs and the transition to a less circular morphology. This is reminiscent of mDia1 KO DCs (24) which were also reported to stay roundish for a prolonged time during cell spreading. The spreading of DCs after seeding on a 2D surface employs the same integrin-and focal adhesion-based mechanisms as in the case of mesenchymal cells. Consistently, mouse embryonic fibroblasts deficient in Ena/VASP proteins did likewise exhibit less efficient cell spreading (12). Since Ena/VASP proteins have close interactions with focal adhesion components, a negative impact of Evl/VASP loss on focal adhesion maturation or dynamics might be the molecular mechanism underlying the observed spreading defect.

The impact of Evl/VASP deficiency on DC migration is complex and dependent on the DC maturation state. For 2D migration on fibronectin, we observe a decrease in speed and directionality for mDCs. This type of DC migration is mechanistically similar to mesenchymal migration, for which the impact of Ena/VASP proteins has already been addressed. While there were conflicting reports about their role in migration in the past (31,32), the most recent rigorous study that evaluated all three family members (12) corroborates our results that Ena/VASP proteins positively regulate migration in 2D.

In the more confined conditions of the under agarose assay, we find again a decrease in speed for iDCs and mDCs, however, directionality is differentially affected. There is no significant difference for iDCs and a slight increase for mDCs. The inherent differences between iDCs and mDCs in their reliance on Ena/VASP proteins become even more evident during 3D migration in collagen gels. In case of iDCs loss of Evl/VASP does not alter cell speed, but substantially increases directionality, while for mDCs cell speed and directionality are both substantially reduced.

The Evl/VASP DKO phenotype in 3D migration is reminiscent of the loss of Hem1, a WRC subunit and thus a stimulator of Arp2/3-dependent actin nucleation. Hem1 KO iDCs show an increase in speed and directionality, while the migration efficiency of Hem1 KO mDCs is significantly impaired (10). Thus, Evl/VASP DKO iDCs resemble Hem1 KO iDCs in their increase in directionality, but in contrast to them they do not migrate with elevated speed. The phenotypic similarity is in line with reports that place VASP and WRC in the same pathway (21,28) by showing for example that VASP enhances WRC stimulation of Arp2/3-dependent actin assembly (21). The phenotypic resemblance in regards to directionality, but not speed, might suggest that the level of WRC activity present in Evl/VASP deficient iDCs is still sufficient to ensure a normal iDC speed. However, to maintain proper protrusion dynamics and thereby limit directionality, an additional stimulation of WRC activity by Ena/VASP proteins might be critical. Also in case of Evl/VASP DKO mDCs it might be uncoordinated protrusion dynamics, which impede efficient directed migration in 3D. However, in Hem1 KO DCs these alterations are tied to obvious cell shape changes. Hem1 KO iDCs migrate faster and more directional due to their pointed shape, while Hem1 KO mDCs display less efficient migration because of their rounded shape with multiple filopodial extensions and reduced protrusion dynamics (10). Evl/VASP DKO DCs do not exhibit such obvious cell shape changes (Figures 1, S3-4). However, it will be important for future studies to examine their protrusion and actin dynamics more closely. The fact that in a previous study with B16-F1 mouse melanoma cells loss of Ena/VASP did not impair invasion in a 3D Matrigel environment (12), is most likely indicative of the distinct migration modes employed by confined DCs and mesenchymal cells.

When we combined Evl/VASP deficiency with different extents of pharmacological formin/myosin or Arp2/3 inhibition, we found far more pronounced effects on cell speed and directionality upon inhibitor treatment than in control cells. Thus, our data show that the presence of Ena/VASP proteins contributes to the known robustness and efficiency of DC migration (6) by rendering DCs more resilient against disturbances in different actin modules. The contribution of Evl and VASP to the overall F-actin level in mDCs is very small and not to be compared with the 50% reduction seen in Hem1 KO mDCs (10). Therefore, it seems likely that the ability of Ena/VASP proteins to interact with the different nucleation machineries is more relevant for shaping local actin dynamics than their own actin nucleation capabilities.

In recent years the coordination and cooperation between the different actin nucleation machineries has in fact gained attention. mDia1 and Arp2/3 were for example shown to cooperatively initiate lamellipodia and ruffles. This cooperation involves the sequential action of both nucleators with mDia providing linear actin filaments for the initial activation of the Arp2/3 complex (33). While this cooperation does not involve direct binding, there are also cases where proteins directly link mDia1 and Arp2/3. For example, Spin90 forms a ternary complex with mDia1 and Arp2/3, which enhances cortical actin generation (34). As discussed, Ena/VASP proteins have already been shown to interact with the WRC in Dictyostelium (28), mammalian cell lines and Drosophila macrophages and to cooperatively enhance Arp2/3 activity (21).

Our proteomic analysis revealed that the known major Ena/VASP family interactions, especially regarding focal adhesion proteins and the WRC, are conserved in iDCs and mDCs. In addition, we retrieved mDia1 as a prominent interactor of the VASP-EVH1 domain thereby extending earlier data about their potential interaction (22) and placing Ena/VASP proteins at the crossroads of the two major actin nucleation pathways that shape DC migration. At the moment it is unclear whether the interactions with WRC and mDia1 occur simultaneously or sequentially. Even though both interactors likely target the same binding site within the EVH1 domain, simultaneous interactions would be conceivable since Ena/VASP proteins multimerize via their EVH2 domain. On the other hand, Arp2/3 functions are rather linked to the cell front where the complex shapes cellular protrusions, while a number of mammalian formins have been shown to rather reside and act at the cell rear (35) including mDia1 which regulates an actin pool at the rear of mature DCs (3) which renders a simultaneous interaction unlikely. In the end live imaging of WRC, mDia1 and Evl/VASP in the different phases of the DC life cycle will be required to unravel their actual spatiotemporal interaction and coordination.

While it would also be highly instructive to unravel the consequence of loss of individual EVH1 interactions, the use of the same binding site renders it very challenging to create binding-deficient mutants for selective interactions. Therefore, at this point the differential consequences of impairing the VASP– mDia and VASP-WRC interaction are still elusive. Nevertheless, we expect the interactions between the three nucleation machineries to contribute to the fine-tuning of the actin network topology for the different requirements of DC physiology.

While our mass spectrometry-based interaction analysis appears highly reliable due to the large number of known and suspected interactors that we recovered, it is of course inherently limited by the fact that we only screened for canonical interactors of the VASP-EVH1 domain that bind via the EVH1 consensus motif. Potential DC-specific interactors of the Pro-rich sequence or EVH2 domain of VASP were not targeted by our approach. In addition, by using recombinant VASP-EVH1 as bait, we were not able to address the potential role of post-translational modifications for modulating DC interactions. Different aspects of Ena/VASP functionality are in fact regulated by phosphorylation at multiple sites (36), and VASP has been shown to be phosphorylated in human DCs upon LPS-induced maturation (37) underlining that Ena/VASP proteins fulfill distinct functions during the DC life cycle.

To unravel whether the actin nucleation capacity of Ena/VASP or their function as interaction platforms are more important for shaping actin-dependent processes of iDCs and mDCs, additional experiments are needed. While we show the importance of Evl and VASP for DC functionality and provide the first characterization of their role in DCs *in vitro*, their exact coordinatory role within the DC actin polymerization network and their physiological relevance for DC-related immune functions await further investigation.

## Materials and Methods

### Animals

The generation of EVL/VASP DKO mice is described in (Kwiatkowski, 2007). Animals were kindly provided by Prof. Frank Gertler (Massachusetts Institute of Technology, USA) via Dr. Stefanie Kliche (Otto-von-Guericke-University, Germany). Animals were housed on a 12 h light/dark cycle with food and water available ad libitum in a specific pathogen-free (SPF) facility. All experiments in the present study were carried out in strict accordance with the guidelines of the Landesamt für Gesundheit und Soziales (LAGeSo) Berlin, and all efforts were made to minimize suffering. Animals were sacrificed by cervical dislocation following isoflurane anesthesia under the permit T0243/08.

Adhering to the 3Rs we reduced animal numbers by switching from breeding EVL^+/−^/VASP^+/−^ mice with each other to a breeding scheme where EVL^+/−^/VASP^+/−^ mice were mated to EVL^-/-^/ VASP^+/−^ mice. This saves animals by increasing the number of DKOs born, but precludes the generation of EVL^+/+^/ VASP^+/+^ animals. Therefore, we used EVL^+/+^/ VASP^+/−^, EVL^+/−^/ VASP^+/+^ and EVL^+/−^/VASP^+/−^ animals as controls in our experiments. We did not observe any differences between these genotypes and EVL^+/+^/ VASP^+/+^ animals.

### Generation of bone marrow-derived dendritic cells

6-20 weeks-old mice were sacrificed by cervical dislocation following isoflurane anesthesia. Abdomen and hind legs were sterilized with 70% ethanol. Murine bone marrow was isolated from the femur and tibiae of the hind legs by flushing the bones with ice-cold, sterile PBS using a 25G needle. Red blood cells were lysed in red blood cell lysis buffer (BioLegend #420301). The resulting cell suspension was centrifuged for 5 min at 500 x g at room temperature. Cells were resuspended in BMDC medium (RPMI 1640 containing 300 mg/l L-glutamine (Gibco #11875-091), supplemented with 10% heat-inactivated fetal bovine serum (Gibco #10270-106), 50 µM β-mercaptoethanol (Gibco #31350-010), 50 U/ml penicillin, 50 U/ml streptomycin (Gibco #15140-163) and 30 ng/ml recombinant mouse GM-CSF (Peprotech #315-03) and seeded into a 15 cm dish. On day 3 GM-CSF-containing RPMI was renewed. On day 6 cells were passaged to two new dishes to dispose of strongly adherent cells. To obtain mature bone marrow-derived DCs, cells in one dish were treated with 0.2 µg/ml LPS (Sigma #0127:B8). Cells in the other dish were left untreated to stay immature. Differentiated immature and mature bone marrow-derived DCs were used for experiments on day 7.

### HEK293T cell transfection with Flag-Mena

70% confluent HEK293T cells were transfected with Flag-tagged Mena in pcDNA3.1+/C-(K)DYK (obtained from Origene; Omu14068c) using calcium phosphate-based transfection. Cells were harvested upon confluency by scraping into PBS after two short PBS washes and processed for immunoblotting as described below.

### Preparation of cell lysates and immunoblot-based analysis

DCs were collected from the plate and centrifuged at 300 x g for 5 min at room temperature. The cell pellet was washed twice with PBS before lysis in lysis buffer (20 mM HEPES pH 7.4, 100 mM KCl, 2 mM MgCl_2_, 1% Triton X-100, 1 mM PMSF, 0.3% mammalian protease inhibitor cocktail (Sigma)). After 30 min on ice lysates were centrifuged at 4°C for 5 min at 17000 x g. The protein concentration of the supernatant was determined by the Bradford assay. Before immunoblotting, lysates were adjusted to 1x Laemmli sample buffer. Samples were analyzed by SDS-PAGE and immunoblotting. Bound primary antibodies (Table 2) were detected by incubation with fluorescently labeled secondary antibodies (IRDuy 800CW or IRDuy 680RD; 1:10.000) using an Odyssey FC Imaging System.

**Table 2:** Antibodies.

| Antigen | Species | Clone name or catalog # | supplier | IF | WB | FACS | comments |
| --- | --- | --- | --- | --- | --- | --- | --- |
| α-tubulin | Mouse | DM1A | Cell signalling | -- | 1:500 | -- | unlabeled |
| β-Actin | Mouse | Ac-15; A5441 | Sigma-Aldrich | -- | 1:1000 | -- | unlabeled |
| CD11c | Hamster | N418; MCD11c05 | Thermo Fisher Scientific | -- | -- | 1:400 | APC-labeled |
| CD40 | Mouse | 3/23; #562846 | BD Biosciences | -- | -- | 1:400 | Pacific Blue-labeled |
| CD86 | Rat | GL1 | eBioscience | -- | -- | 1:100 | APC-labeled |
| EVL | Rabbit | TA343847 | Origene | -- | 1:500 | -- | unlabeled |
| GAPDH | Mouse | G8795 | Sigma | -- | 1:1000 | -- | unlabeled |
| mDia1 | Rabbit | Ab129167; Ab96784 | Abcam | -- | 1:1000 | -- | unlabeled |
| mDia1 | Mouse | Clone 51; 610848 | BD Biosciences | 1:50 | -- | -- | unlabeled |
| Lamellipodin | rabbit | -- | Matthias Krause, Kings College London | -- | 1:20000 | -- | unlabeled |
| Mena | Rabbit | LS-C170270 | LSBio | -- | 1:500 | -- | unlabeled |
| p34/ARPC2 | Rabbit | 07-227-I | Sigma-Aldrich | 1:100 | 1:1000 | -- | unlabeled |
| Paxillin | Rabbit | Y113; ab32084 | Abcam | 1:400 | -- | -- | unlabeled |
| VASP | Rabbit | 9A2 | Cell signalling | -- | 1:500 | -- | unlabeled |
| Vinculin | Mouse | V9264 | Sigma | -- | 1:1000 | -- | unlabeled |
| Vinculin | Rabbit | Ab73412 | Abcam | 1:400 | -- | -- | unlabeled |
| Wave2 | Rabbit | D2C8 | Cell signalling | -- | 1:500 – 1:1000 | -- | unlabeled |
| Wave2 | Mouse | C-6; sc-373889 | Santa Cruz Biotechnology | 1:100 | -- | -- | unlabeled |
| Zyxin | Rabbit | Z4751 | Sigma | 1:400 | 1:500 | -- | unlabeled |

### Flow cytometry-based assays

Flow cytometry was used to quantify protein surface levels, polymerized F-actin and macropinocytosis. For surface stainings of differentiation and maturation markers, PBS-washed cells were pelleted by centrifugation at 500 x g for 5 min. Pellets were resuspended in PBS containing 10% FBS and the respective antibody (s. Table 2 for concentrations). Cells were incubated on ice protected from light for 20 min before 2x washing in PBS containing 10% FBS. Samples were immediately processed with a BD LSR Fortessa (BD Biosciences) using BD FACS Diva™, and the data were analyzed using FlowJo.

For the detection of polymerized actin by intracellular phalloidin staining, DCs were harvested and washed with PBS. Cells were fixed with 4% PFA/4% sucrose for 15 min at room temperature. After washing with PBS cells were permeabilized by a 30 min incubation in 0.05% Triton-X100, 2% BSA in PBS at room temperature. After washing with PBS cells were incubated with 1:1000 Phalloidin iFluor 594 (Abcam, ab176757) for 1 h on ice in the dark. Cells were washed again with PBS and immediately processed with a BD LSR Fortessa (BD Biosciences) using BD FACS Diva™. The data were analyzed using FlowJo.

For quantifying macropinocytosis immature and mature DCs were washed with PBS, resuspended in a complete culture medium containing 1 mg/ml FITC-dextran (70,000 MW FITC-dextran; Thermo Fisher Scientific), and incubated at 37°C and 5% CO_2_. The cells meant as a negative control for unspecific surface binding were incubated with FITC-dextran at 4°C. After 1 h the uptake was stopped by two washes with ice-cold FACS buffer (0.1% sodium azide, 2% FBS, 10 mM EDTA in PBS pH 7.5), and samples were rapidly processed with a BD LSR Fortessa (BD Biosciences) using BD FACS Diva™. The data were analyzed using FlowJo.

### Immunofluorescence

Mature or immature DCs were harvested, seeded onto 50 µg/ml fibronectin-coated coverslips, and allowed to adhere for 4 h. For immunofluorescence stainings, cells were fixed with 4% PFA in PBS for 10 min at room temperature. Cells were permeabilized and blocked in goat serum dilution buffer (GSDB) containing 0.3% Triton X-100 and 25% goat serum in PBS for 30 min. Afterwards, samples were incubated with different primary antibodies in GSDB for 20 h at 4°C. After three short PBS washes, coverslips were incubated with species-specific Alexa488- or Alexa647-labeled secondary antibodies (Invitrogen) at a 1:200 dilution in GSDB for 1 h in the dark at room temperature. To visualize F-actin, Alexa568-coupled phalloidin (Invitrogen) was applied at a 1:50 dilution in GSDB together with the secondary antibodies. After the incubation, coverslips were washed three times with PBS and mounted onto microscopy slides with ImmuMount mounting solution (Thermo Fisher Scientific) supplemented with 1 μg/ml DAPI to stain nuclei. Images were taken by spinning disc confocal microscopy using a Nikon Eclipse Ti microscope operated by NIS-Elements (Nikon), equipped with an AU-888 EMCCD camera (Andor) and a CSU-X1 spinning disc (Yokogawa). Image analysis was performed with Fiji, an ImageJ 1.53c package.

### Spreading assay

Mature DCs were harvested, washed with PBS, and seeded on glass coverslips that were coated with 50 µg/ml fibronectin. Cells were allowed to spread for 10, 20, 30, 40 or 60 min before fixation with 4% PFA in PBS for 10 min at room temperature. After 2 washes with PBS cells were permeabilized and blocked with 0.3% Triton-X100 and 10% fish gelatin in PBS for 5 min. After 3 washes with PBS cells were incubated with 1:50 Phalloidin-Alexa595 for 1 h at room temperature in the dark. After 3 washes with PBS coverslips were mounted onto microscopy slides with ImmuMount mounting solution. Images were acquired on a confocal microscope and cell area and circularity were quantified using ImageJ software.

### 2D chemotaxis assay

The migration chamber, tracking, and quantification were performed as described in detail in (38) with the following modifications: The migration chamber was coated with 10 µg/ml fibronectin for 1 h at 37°C using 150 µl of solution. Afterward, the fibronectin solution was aspirated, and the chamber was washed with 150 µl PBS. After removal of the PBS 12.500 mature DCs in 150 µl full medium were transferred into the migration chamber and left to adhere for 1-2 h at 37°C and 5% CO_2_. Once cells adhered to the chamber, 230 µl of medium were gently added and the migration chamber was carried to the microscope. Right before imaging 5 µl of 550 ng/ml CCL19 were added to the medium in the migration chamber. Imaging was performed on a Nikon Eclipse Ti epifluorescent microscope, equipped with an Andor sCMOS camera, an Okolab incubator for life cell imaging (set to 37°C and 5% CO_2_) using a 4x objective and 1.5x added zoom and PhL phase contrast. Images were acquired every 35 s for 6 h.

### Under agarose assay

The under agarose migration analysis was performed as described in (8,39) using 35 mm glass-bottom dishes (Ibidi #81158). The day before the migration assay the 35 mm dishes were incubated with 2 ml of 20% FBS solution at 4°C overnight. The next day the FBS solution was removed and the dish briefly washed with PBS. 2 ml of 56°C warm gel solution (1 volume RPMI/20% FBS, 1 volume 2x HBSS, 2 volumes 2.5% agarose in water (ultrapure agarose, Invitrogen, #16500-100)) were cast per 35 mm dish and allowed to solidify. After the gel solidified, two holes were punched into the agarose (3 mm wide with 3 mm distance between the hole rims), before it was incubated for 30-45 min at 37°C and 5% CO_2_. One hole was filled with 50 µl medium containing 1.2 µg/ml CCL19. The second hole was filled with 50 µl medium containing 5x 10^5^ cells. 2.5 – 3 h later, when cells had come into the field of view between the holes, imaging was started on a Nikon Eclipse Ti epifluorescent microscope, equipped with an Andor sCMOS camera, an Okolab incubator for life cell imaging (set to 37°C and 5% CO_2_) using a 10x objective with phase contrast. Images were acquired every 1 min for 6 h. For quantification, cells were tracked manually by MtrackJ plug-in in Fiji as described in (40). Further analysis was performed with the help of the DiPer suite as described in (25).

### 3D chemotaxis assay

The migration chamber, tracking, and quantification were performed as described in detail in (38) with the following modifications: The chosen collagen gel concentration was 1.9 mg/ml. The chosen CCL19 concentration was 650 ng/ml. The CCL19 was mixed into the medium on top of the gel to create a chemokine gradient. Imaging was performed on a Nikon Eclipse Ti epifluorescent microscope, equipped with an Andor sCMOS camera, an Okolab incubator for life cell imaging (set to 37°C and 5% CO_2_) using a 4x objective and 1.5x added zoom and PhL phase contrast. Images were acquired every 2 min for 6 h. For evaluating the effect of the formin inhibitor SMIFH2 the compound was added to a final concentration of either 5 µM or 10 µM to the collagen gel together with 14000 cells and also to the 200 µl of CCL19 solution that was filled on top of the solidified gel to form the chemokine gradient.

### Pulldown assay

A construct encoding GST-tagged VASP-EVH1 domain (Swissprot ID P50552; amino acids 1-113; vector backbone pGEX-4T1) was provided by the group of Dr. Ronald Kühne. GST and GST-VASP-EVH1 were purified from overexpressing E. coli using GST-bind resin (Novagen) according to standard protocols. Beads were incubated with immature and mature DC lysates prepared in RIPA buffer (10 mM HEPES, 50 mM NaCl, 1% NP-40, 0.5% DOC, 0.1% SDS, protease inhibitor cocktail (Roche), 1 mM Na_3_VO_4_,1 mM DTT, benzonase, pH 7.7). To some samples, 100 µM of the EVH1-specific inhibitor ProM (27) was added (Note: The used ProM ligand version did not efficiently cross DC cell membranes in our culture conditions, thus it could be used only on cell lysates, not for living cells.). After the incubation, beads were thoroughly washed with RIPA buffer without protease inhibitor cocktail, Na_3_VO_4_, DTT and benzonase. After elution interacting proteins were initially identified by mass spectrometry. Subsequently, a set of interactors was verified by independent pulldown experiments that were analyzed by SDS-PAGE and immunoblotting.

### Mass spectrometry

#### Experimental Design

Two independent pull-down experiments per sample were performed (n=2; based on prior experience with expected variability) with either immature or mature DC lysates (untreated or incubated with 100 µM of the EVH1-specific inhibitor ProM (27)).

#### Sample preparation

Proteins were loaded onto SDS-PAGE and subjected to in-gel digestion. In brief, gel bands were reduced with 5 mM DTT at 56°C for 30 min and alkylated with 40 mM chloroacetamide at room temperature for 30 min in the dark. Protein digestion was carried out using trypsin at an enzyme-to-protein ratio of 1:20 (w/w) at 37°C overnight.

#### LC/MS and data analysis

LC/MS analysis was performed using an UltiMate 3000 RSLC nano LC system coupled on-line to an Orbitrap Elite mass spectrometer (Thermo Fisher Scientific). Reversed-phase separation was performed using a 50 cm analytical column (in-house packed with Poroshell 120 EC-C18, 2.7µm, Agilent Technologies) with a 120 min gradient. Label-free quantification was performed using MaxQuant (version 1.6.1.0) using the following parameters: MS ion mass tolerance: 4.5 ppm; MS2 ion mass tolerance: 0.5 Da; variable modification: Cys carbamidomethyl, Cys Propionamide, Met oxidation; protease: Trypsin (R,K); allowed number of mis-cleavage: 2; database: SwissProt database of Mouse (2016 oktuniprot-proteome, number of sequences: 22136); label free quantification and match between runs were enabled. Results were reported at 1% FDR at the protein level.

#### Analysis of obtained results based on LFQ intensity values

The results obtained with immature and mature DCs (iDC and mDC) were analyzed separately. In each experiment, biological replicates were performed and two conditions (i.e., inhibitor treated and untreated cells) were compared. Of the ~300 detected proteins we excluded all where no protein was detected in the untreated condition in one or both of the two experiments. For the remaining hits, we calculated the ratio between the LFQ intensity of the untreated condition and the EVH1-inhibitor treated condition since specific EVH1 domain interactors of VASP should be competed off by the inhibitor. We included proteins as hits that had a log2 ratio of intensities (untreated over inhibitor treated) > 2.5 in both experiments. In addition, we included the proteins detected in the untreated, but not detected in the EVH1-inhibitor treated sample as potential candidates.

### Statistical analyses

For data transparency values are depicted as individual dots and also as mean±SEM (standard error of the mean). Statistical significance of data was analyzed by the statistical tests indicated in the figure legends using GraphPad Prism 9.1 software. The level of significance is indicated in the following way: ****=p<0.0001; ***=p<0.001; **=p<0.01; *=p<0.05. Data with arbitrary absolute values, like fluorescent intensities, which for technical reasons showed high variability between experiments, was normalized before performing statistics. The number of analyzed cells and independent experiments is stated in each figure legend.

## Supporting information

This article contains supporting information.

## Acknowledgements

We would like to thank the Imaging Facility and the Mass Spectrometry facility of the FMP for continuous support, especially Heike Stephanowitz for processing MS samples. We thank Claudia Schmidt for expert technical assistance with genotyping, DC culturing, and immunoblotting. We are indebted to Frank Gertler for generously sharing his mouse line. We would also like to thank Matthias Müller and Matthias Barone for sharing their expertise with the EVH1 inhibitor and Volker Haucke for stimulating discussions.

## Author contributions

S.P.V. developed the project, established and performed cell migration assays and biochemical experiments. H.N. carried out the cell spreading and macropinocytosis assays. L.H. contributed immunofluorescence analyses. M.G. performed the differentiation and maturation analyses. F.L. supervised mass spectrometry. R.K. contributed valuable reagents and advice. T.M. designed and supervised the project and wrote the manuscript with input from all authors.

## Funding

This work has been supported by an Integrated Project Grant from the FMP to T.Maritzen and R. Kühne and by a grant from the Deutsche Forschungsgemeinschaft (DFG, German Research Foundation) to T.M. (461336323).

## Conflict of interest

The authors declare that they have no conflicts of interest with the contents of this article.

## Notes

### Competing Interest Statement

The authors have declared no competing interest.

