## Supplementary Figures for "Ena/VASP proteins at the crossroads of actin nucleation pathways in dendritic cell migration"

**Supplementary Figure S1**
Analysis of Ena/VASP family protein expression in control and Evl/VASP DKO DCs

**Supplementary Figure S2**
DC Differentiation and maturation are unaltered in absence of Evl and VASP

**Supplementary Figure S3**
Localization and levels of focal adhesion and actin regulatory proteins in Evl/VASP DKO iDCs

**Supplementary Figure S4**
Localization and levels of focal adhesion and actin regulatory proteins in Evl/VASP DKO mDCs

**Supplementary Figure S5**
Immunoblot analysis of focal adhesion and actin regulatory proteins in Evl/VASP DKO DCs

**Supplementary Figure S6**
Double deficiency of Evl and VASP is required to cause changes in 3D migration

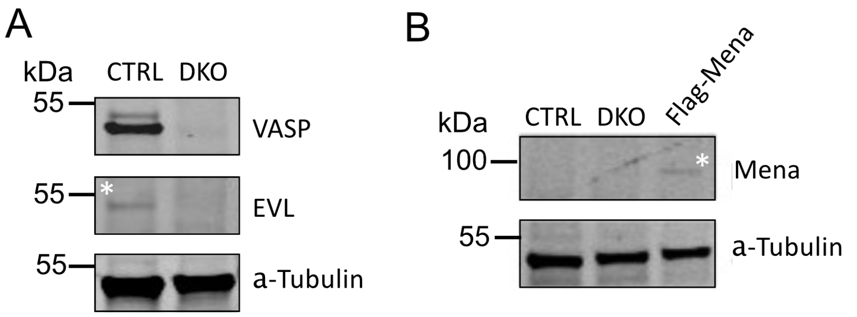


**Supplementary Figure S1**
**Analysis of Ena/VASP family protein expression in control and Evl/VASP DKO DCs**
(A) VASP and Evl can be detected in control DCs and are absent in DKO DCs. Lysates of mature WT (labeled CTRL) and Evl/VASP DKO DCs were analyzed by immunoblotting using the indicated antibodies. Detection of α-Tubulin was used as loading control. VASP could easily be detected in CTRL DCs, while Evl only gave a faint signal. (B) Since Mena could not be detected at all in DCs, HEK293 cells transfected with Flag-Mena were used to demonstrate that the antibody is in principle able to detect Mena.

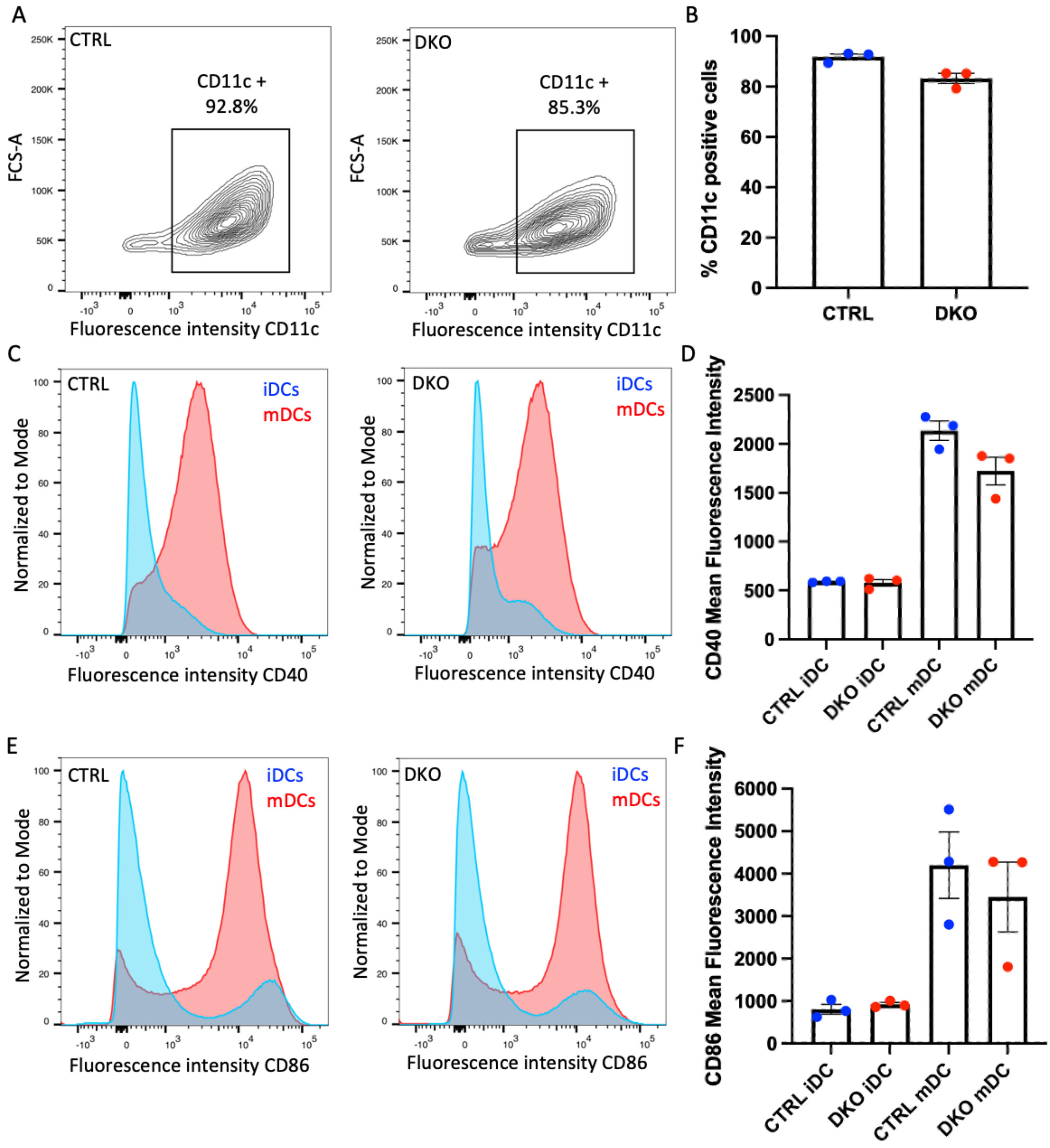


**Supplementary Figure S2**

**DC Differentiation and maturation are unaltered in absence of Evl and VASP**

Bone marrow-derived control and DKO DCs both display classical markers of DC differentiation and maturation. (A,B) Control and DKO DCs were analyzed by flow cytometry with CD11c-specific fluorescently labeled antibodies. The plots depict the set gate and the % of CD11c positive cells for an exemplary experiment. Cells with a fluorescence intensity of at least $10^3$ were defined as CD11c-positive amounting to 92.8% for the depicted control and 85.3% for the DKO DC culture. The bar diagram shows the % of CD11c positive cells for 3 independent experiments. (C-F) Immature control and DKO DCs were left untreated (blue line) or were matured by incubation with LPS for 24 h (red line) and subsequently analyzed by flow cytometry with fluorescently labeled antibodies against the DC maturation markers CD40 (C-D) or CD86 (E,F). (C,E) The graphs depict the normalized number of cells counted at different fluorescence intensities for both conditions illustrating the increase in surface expression of the tested markers after LPS treatment. (D,F) Quantification of mean fluorescence intensity for the indicated marker proteins in immature and LPS-matured control and DKO DCs based on flow cytometry (N=3 independent experiments).

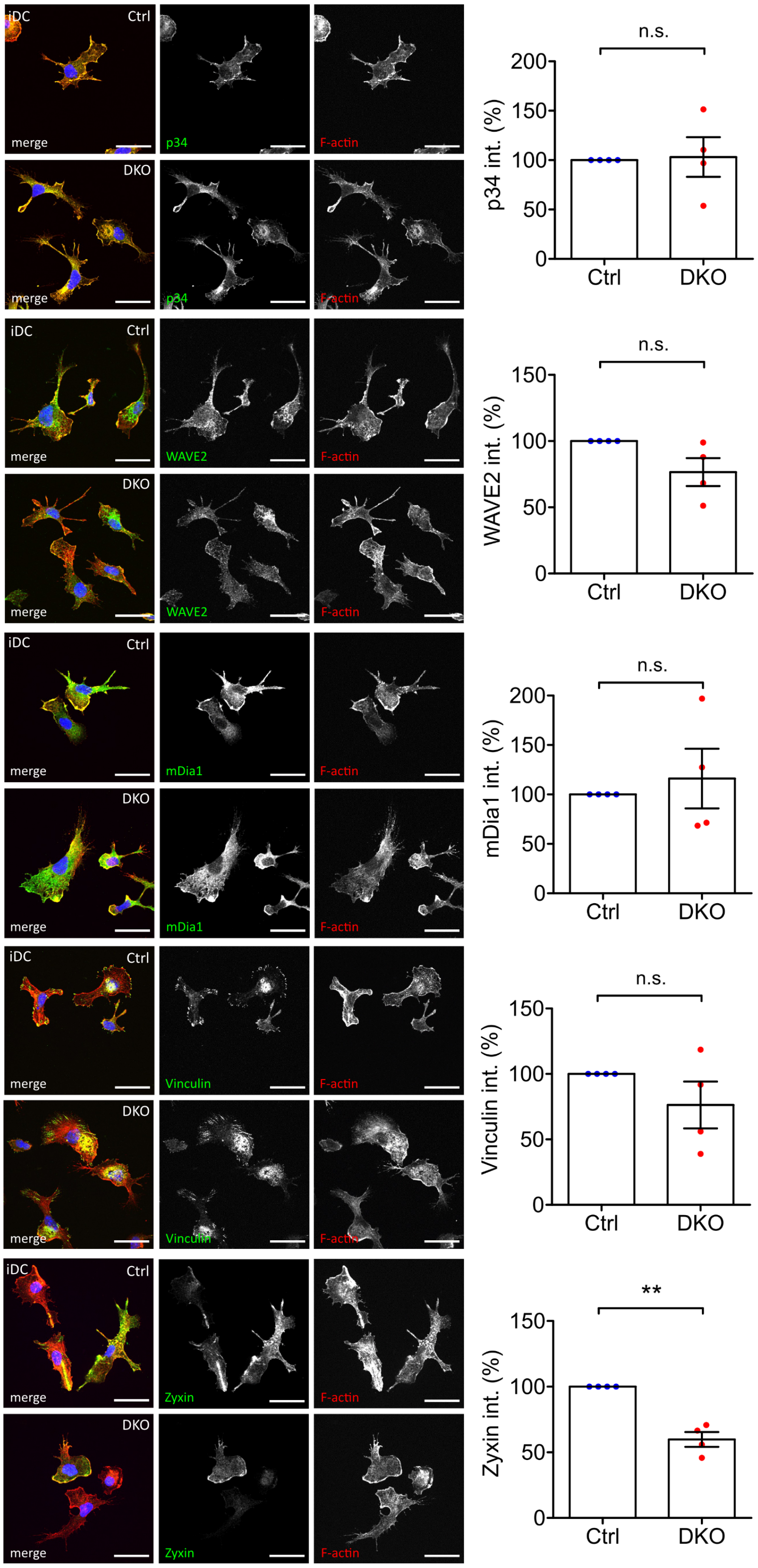


**Supplementary Figure S3**

**Localization and levels of focal adhesion and actin regulatory proteins in Evl/VASP DKO iDCs**

Levels and localization of focal adhesion and actin regulatory proteins are mostly unaltered in DKO iDCs. Immature control and Evl/VASP DKO DCs were processed for immunofluorescence and labeled with the indicated antibodies. Left: Images representative of 4 independent experiments. Nuclei were stained with DAPI and are depicted in blue in the merged images. Scale bar: 25 µm. Right: Quantification of fluorescence intensities which are expressed as % of control levels (depicted as mean±SEM; statistical analysis by One-sample t-test; N=4 independent experiments; ns=non significant; **p<0.01).

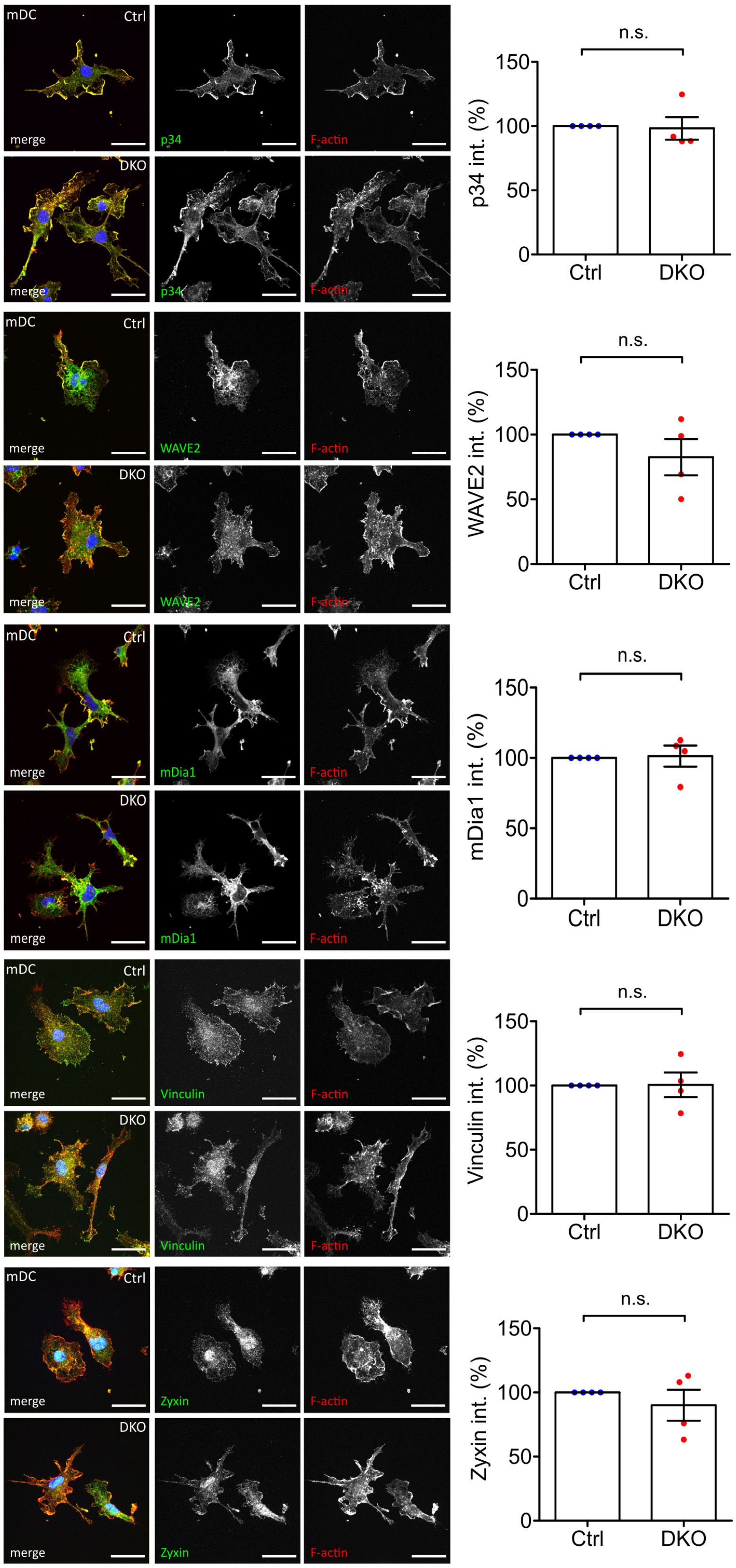


**Supplementary Figure S4**

**Localization and levels of focal adhesion and actin regulatory proteins in Evl/VASP DKO mDCs**

Levels and localization of focal adhesion and actin regulatory proteins are unaltered in DKO mDCs. Mature control and Evl/VASP DKO DCs were processed for immunofluorescence and labeled with the indicated antibodies. Left: Images representative of 4 independent experiments. Nuclei were stained with DAPI and are depicted in blue in the merged images. Scale bar: 25 µm. Right: Quantification of fluorescence intensities which are expressed as % of control levels (depicted as mean±SEM; statistical analysis by One-sample t-test; N=4 independent experiments; ns=non significant; **p<0.01).

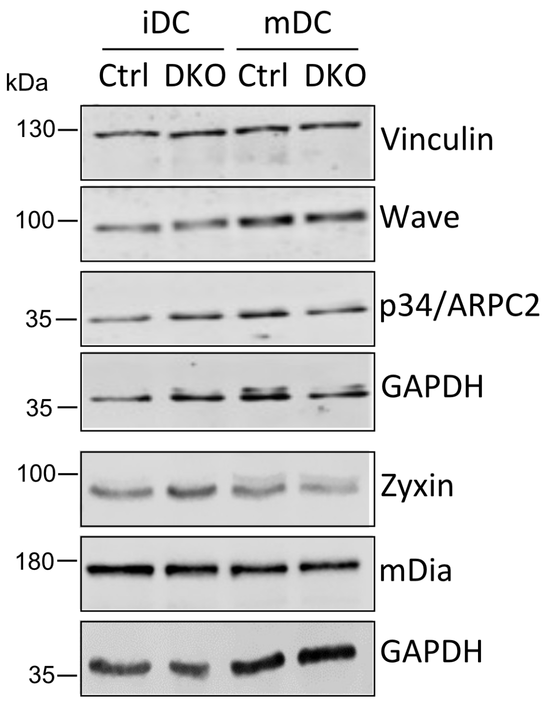


**Supplementary Figure S5**
**Immunoblot analysis of focal adhesion and actin regulatory proteins in Evl/VASP DKO DCs**
No change in overall protein levels of focal adhesion and actin regulatory proteins. Lysates of immature and mature control and DKO DCs were probed with the indicated antibodies.

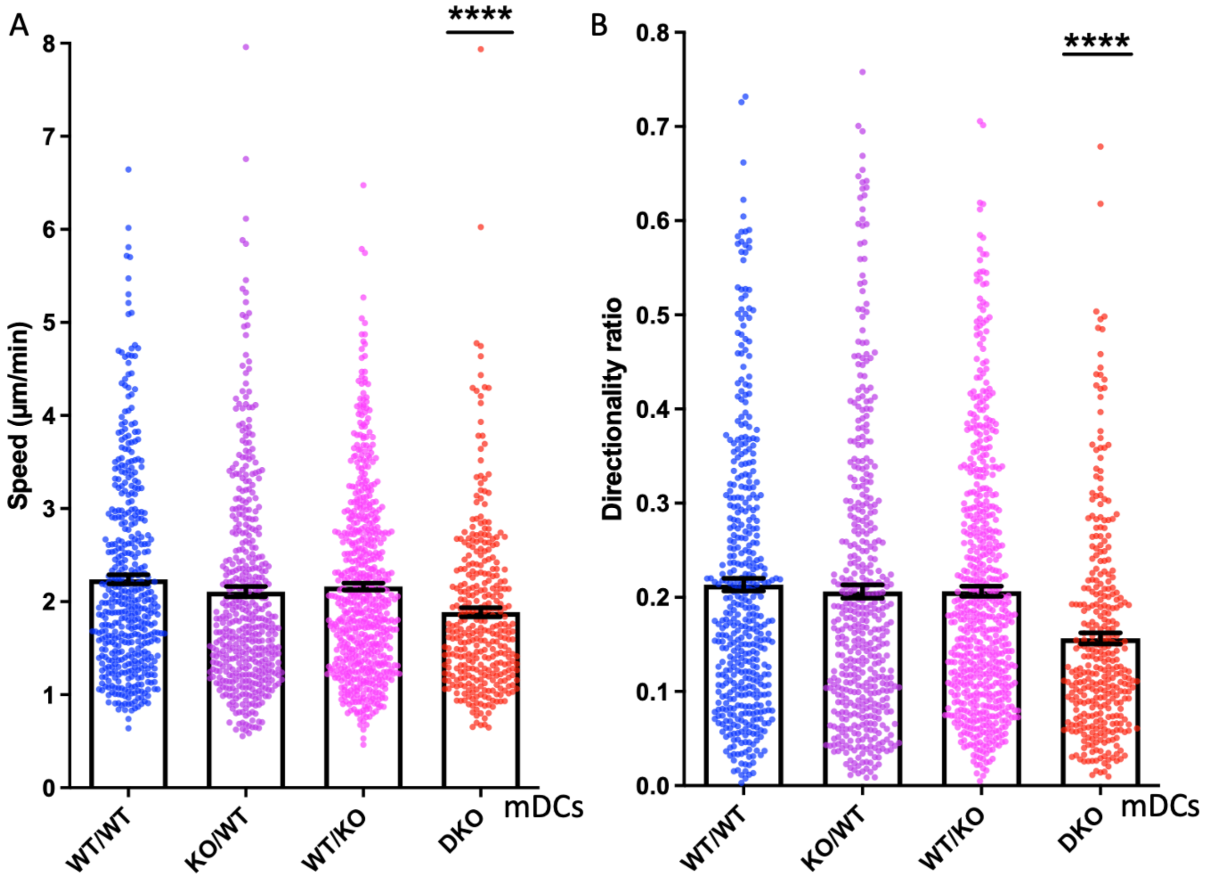


**Supplementary Figure S6**

**Double deficiency of EVL and VASP is required to cause changes in 3D migration**

Mature $EVL^{wt}/VASP^{wt}$ (WT/WT), $EVL^{ko}/VASP^{wt}$ (KO/WT), $EVL^{wt}/VASP^{ko}$ (WT/KO) and $EVL^{ko}/VASP^{ko}$ (DKO) DCs were embedded into 1.9 mg/ml collagen gels to compare the effect of single and double loss of EVL and VASP. The upper gel surface was covered with medium containing 650 ng/ml CCL19. DC migration was monitored by bright-field real-time microscopy for 6 h. Images were obtained at 2 min intervals and analyzed in automated manner. Speed (A) and directionality (B) are depicted as mean±SEM (N(WT/WT)=455, N(KO/WT)=468, N(WT/KO)=629, N(DKO)=341 cells from 3 (for all except DKO) respectively 2 (for DKO) independent experiments; statistical analysis by Kruskal-Wallis test followed by Dunn´s multiple comparison test; ****p<0.0001).
